# Single-nucleus transcriptomics links telencephalic cell signatures to ocean acidification effects on social behaviour in a coral reef fish

**DOI:** 10.64898/2026.10.28.762664

**Authors:** Debora Desantis, Celia Schunter

## Abstract

Mutualistic interactions are fundamental to the health and stability of reef ecosystems. While elevated CO_2_ has been implicated in impairments in interaction behaviours, how this environmental change alters cellular states and intercellular communication in the brain remains unresolved. Here, by analysing the single-nucleus RNA profile of ∼66,000 telencephalic nuclei from a keystone mutualist, the cleaner wrasse (*Labroides dimidiatus*), interacting under ambient or elevated CO_2_ conditions, we identify highly cell type specific neural responses to elevated CO_2_. We highlight a substantial depletion in Oligodendrocyte Precursor Cells (OPCs), notably linked to oligodendrocyte myelination and neural structure maintenance. Elevated CO2 also elicited targeted transcriptional shifts across discrete cell types, characterised by widespread glutamatergic signalling and altered co-expression networks associated with pH regulation, energetic metabolism, and circadian rhythm, thereby compromising the cleaner’s physiologic equilibrium. Additionally, cell-cell communication was altered under elevated CO_2_, including remodelling in SEMA4 and PTPR signalling pathways within radial glial and OPCs, respectively, revealing network signalling rewiring in response to the changed environment. These findings uncover the high-resolution neural substrates vulnerable to ocean acidification, providing mechanistic insight into how climate-driven environmental changes alter brain function and destabilise complex mutualistic ecological interactions.

## INTRODUCTION

Ocean chemistry is rapidly changing as the oceans absorb a substantial fraction of anthropogenic CO_2_ emissions. The resulting increase in proton concentration ([H^+^]) reduces seawater pH, a process known as ocean acidification (OA), which is predicted to further intensify under future climate-change scenarios ^1–3^. While elevated CO2 has been implicated in altering aspects of fish behaviour, cognition, circadian regulation and sensory functions ^4–17^, the literature increasingly reveals that these responses are complex and highly variable both within and among species ^8,18–23^. Rather than a single uniform impairment, this pronounced inter- and intra-individual variation highlights a critical knowledge gap: macroscopic behavioural observations alone are insufficient to resolve the hidden neurological impacts of environmental stress. Consequently, fine-scale, high-resolution cellular analyses are urgently needed to disentangle exactly how the brain responds to shifting CO_2_ levels. Uncovering these molecular vulnerabilities is crucial for understanding the broader ecological implications, as complex inter- and intraspecific social interactions are fundamental to the structure and functioning of marine communities ^24–26^.

A keystone example of such dynamics in the marine ecosystem is the mutualism between the coral reef Bluestreak cleaner wrasse (*Labroides dimidiatus)* (i.e. cleaner) and its client fish. By providing cleaning services ^27^, cleaners monitor ectoparasite load ^27,28^, shape reef fish diversity and community structure ^26,29–31^ and even enhance client cognitive health and cognitive performance ^32^. To handle this diverse behavioural network cleaners employ sophisticated social and cognitive abilities, comparable to those of some primates ^33,34^, enabling them to form and maintain long-term relationships that scale up to influence entire reef communities ^26^. However, environmental perturbations like OA threaten to destabilise these delicate interaction networks. Studies on both laboratory-acclimated and wild populations indicate that elevated CO_2_ can alter cleaners’ motivation to interact and decrease the quality of the service they provide ^11,12,14,35^. Furthermore, near-future CO_2_ conditions have been linked to shifts in cleaner’s cognitive performances ^12^, including processes essential for client recognition ^36^. Because cleaner behaviour directly shapes local fish communities ^26^, these individual-level behavioural shifts have the potential to propagate, posing a severe risk to broader reef ecosystem functioning. Yet, despite the clear ecological interest, the underlying neural substrates and specific cellular vulnerabilities that govern these behavioural responses to environmental stress remain largely unexplored.

Initial investigations into the neural mechanisms underlying these behavioural shifts have linked OA to altered transcriptional regulation within specific brain regions of the cleaner wrasse ^14^. Previous bulk transcriptomic and physiological analyses suggest OA affects broad neuromodulatory systems, including glutamatergic, GABAergic, dopaminergic and serotonergic pathways, as well as networks related to learning, memory and visual perception ^11,14^. The forebrain and particularly, the telencephalon, is of specific interest due to its central role in regulating social behaviour across vertebrates ^37^, raising concerns regarding how OA perturbs this crucial brain region.

To date, much of the mechanistic focus in OA research has centred on the hypothesis that altered GABAergic neurotransmission drives behavioural disruptions ^38–40^. While this mechanism is supported by various behavioural studies ^4,7,9,35,41–43^, complex social behaviours, are regulated by multiple neuromodulatory systems. For example, dopaminergic and serotonergic systems dictate the motivation of cleaner fish to interact ^44–48^, both of which are disrupted under OA ^11^. Because GABAergic circuits also modulate serotonergic and dopaminergic responses ^49,50^, a complex network of neurotransmitter crosstalk likely underpins the brain’s response to OA. Furthermore, immediate early genes (IEGs), which are critical markers of neuronal activation, synaptic plasticity, and memory consolidation ^51^, are involved in cleaner-client interactions ^47^ and may represent primary responders to altered CO_2_ ^8^. Crucially, however, all prior transcriptomic studies on OA have relied on bulk tissue analyses. By averaging signals across highly heterogeneous cell populations, bulk RNAseq obscures the precise cellular targets of environmental changes, leaving the true high-resolution mechanisms unresolved.

Here, we combine single-nucleus RNA sequencing (snRNA-seq) with behavioural analyses to identify the specific cellular substrates of OA-induced behavioural alterations within the telencephalon of *L. dimidiatus*. By comparing fish interacting under present-day and near-future OA conditions, we characterise shifts in cellular composition, transcriptional profiles, gene co-expression networks and cell-cell communication. Given the established roles of dopaminergic, serotonergic, and GABAergic signalling in modulating mutualistic interaction behaviour in cleaners ^44–46,48–50^, we hypothesise that OA would alter the relative abundance and gene expression profiles of specific neuromodulatory cell population. By mapping these responses at a single-cell resolution, we evaluate how OA remodels cellular communication states across neuronal and glial populations. Ultimately, our study identifies the precise neuronal cell types most vulnerable to near-future OA, providing unprecedented mechanistic insight into how climate-driven changes in ocean chemistry may destabilise social behaviour and complex mutualistic interactions.

## RESULTS

### Single nuclei transcriptional profile of the telencephalon reflects conserved cell classes

To characterise the cellular heterogeneity in the telencephalon of the cleaner wrasse, we performed snRNA-seq using nuclei dissociated from a total of six individuals, three per condition, generating 66,878 high-quality nuclei (Fig. 1a; Figs. S1 and S2; Supplementary Excel Table S1a,b). We identified 33 distinct cell clusters, with nuclei from all samples contributing to the configuration of the integrated clusters (Fig. S3a,b) and confirming their diversity by plotting the top five marker genes for each identified cluster (Fig. S4; Supplementary Excel Table S2). Next, we classified the identified cell clusters into major neuronal and non-neuronal classes (Fig. 1b,c) according to the expression profile of cell type specific marker genes identified from the literature (Fig. 1d; Supplementary Excel Table S3). Based on their expression profile, neurons were classified as either GABAergic (*gadb1*^+^, *gad2*^+^, *slc32a1*^+^), or glutamatergic (*slc17a6*^+^, *adcyap1b*^+^), which represented the most abundant cell type identified (42,636, comprising 63%) (Supplementary Excel Table S4). Among the non-neuronal cells, we identified clusters expressing characteristic markers of oligodendrocytes (*mbp*^+^, *mpz*^+^, *olig2*^+^, *myrf*^+^, *ninj2*), oligodendrocyte precursor cells (i.e. OPCs; *olig2*^+^, *cspg4*^+^), and radial glial cells (i.e. RGc; *gfap*^+^, *cyp19a1b*^+^, *slc1a2b*^+^). We also assigned a small population of cells expressing lymphatic endothelial markers to putative perithelial cells (Perithelial; *vegfd*^+^) ^52^. Lastly, one cluster could only be categorised at the pallial level, as it exhibited pallial features but did not express any specific canonical marker genes (Fig. 1c,d, Supplementary Excel Table S3). Cell type transcriptional relationships were further supported by unbiased tree reconstruction, in which neuronal clusters were clearly defined and separated from non-neuronal clusters, with high bootstrap support reinforcing the cluster subdivision (Fig. 1c).

**Fig. 1.**
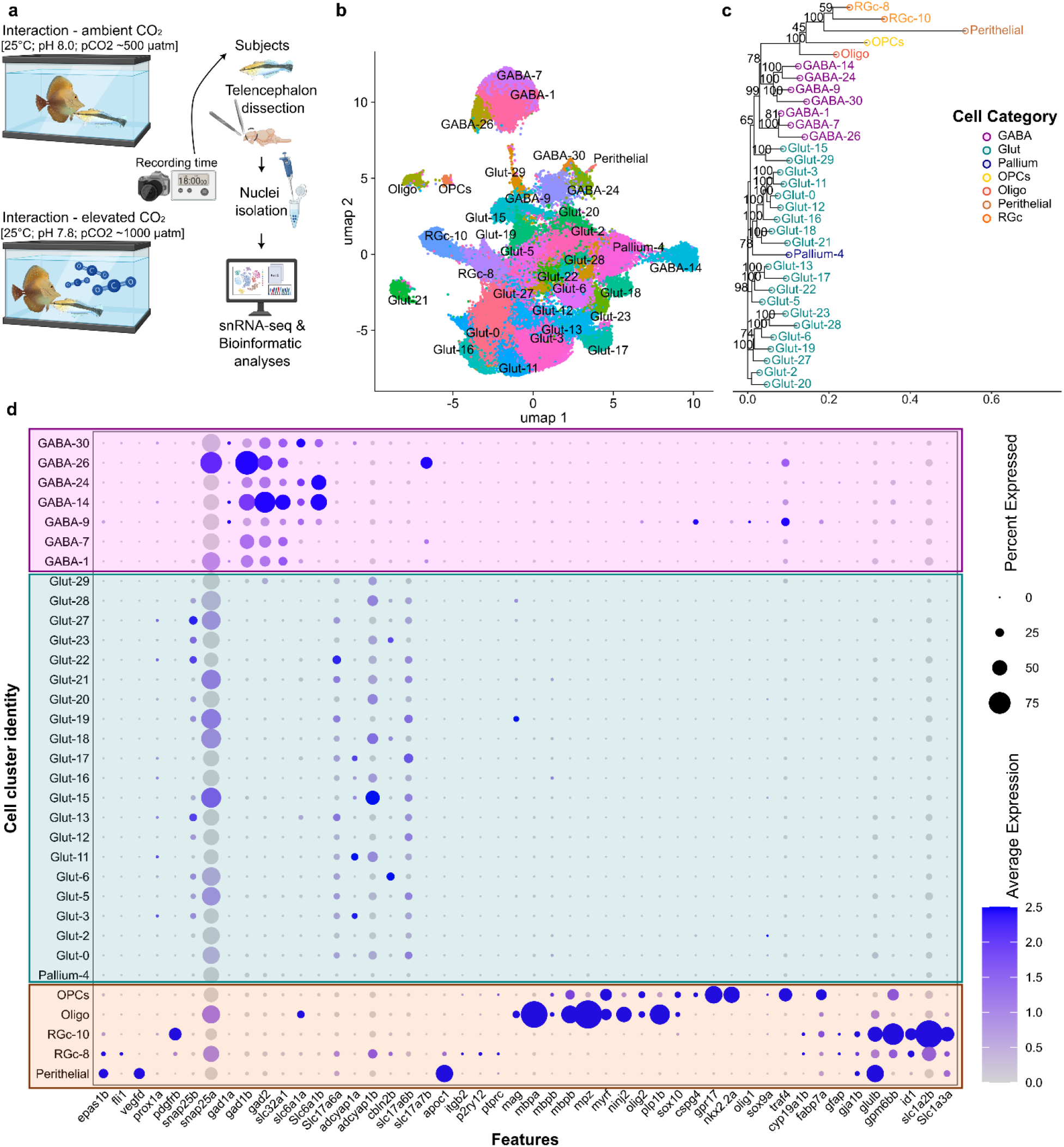
Experimental design and cellular profile of *L. dimidiatus* telencephalon. **(a)** Behavioural trial where *Labroides dimidiatus* (cleaner, on the right side of the tank) and *Zebrasoma scopas* (client, on the left side of the tank) were allowed to interact in present day scenario CO_2_ (ambient CO_2_, cleaners *n = 12;* clients *n =6*) or future elevated CO_2_ conditions (elevated CO_2_ treatment, cleaners *n = 12;* clients *n =6*) for 18 minutes following telencephalon dissection, nuclei isolation, sequencing and bioinformatics analyses (*n = 3* subjects per each treatment). **(b)** Integrated UMAP with cell clusters (*n = 33*) assigned. **(c)** Cell type tree for *L. dimidiatus* reconstructed by using all expressed genes (10,000 bootstraps) **(d)** Dotplot representing the percentage of nuclei and the average expression of selected marker genes employed to assign broad cluster identities. GABAergic clusters (purple), Glutamatergic/Pallial clusters (cyan), non-neuronal clusters (orange). See Supplementary Excel Table S3 for more information on the marker gene list.

### Distinct molecular signalling systems emerged from the identified cell clusters

To uncover neuronal signatures and identify transcriptional programs associated with behaviourally relevant functions, we further investigated the co-expression of genes specifically related to neuromodulatory systems and social behaviour in fish. To this end, we generated a curated gene list comprising genes encoding neuropeptides as well as components of small-molecule neurotransmitter pathways (Fig. 2a; Supplementary Excel Table S5).

**Fig. 2.**
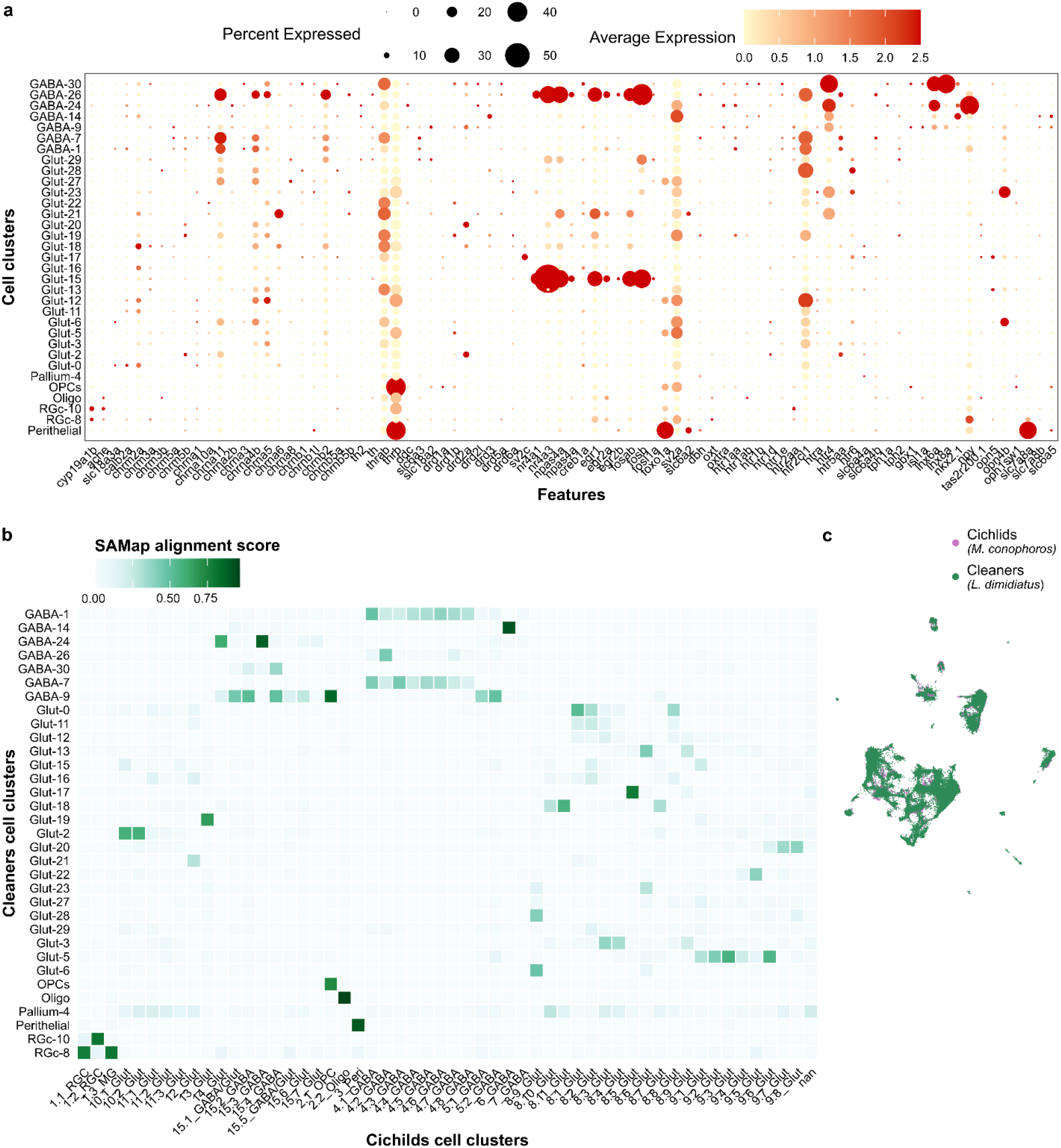
Molecular signatures and neuroanatomical architecture of identified cell clusters. **(a**) Dotplot representing the percentage of nuclei and the average expression level of selected genes involved in neuromodulatory system and social behaviour in the *L. dimidiatus* (see Supplementary Excel Table S5 for more information on the reference gene list. **(b)** Cell cluster mapping comparisons between snRNA-seq cleaner fish (*L. dimidiatus*) and cichlid (*Mchenga conophoros*) dataset. Darker colours indicate higher SAMap mapping scores. See Supplementary Excel Table S6b for the complete list of SAMap alignment scores. **(c)** SAMap projections within a UMAP space showing the telencephalic cleaner (*L. dimidiatus*) nuclei (green, n = 66,878) and cichlid (*M. conophoros*) nuclei (purple, n = 33,895). Cleaner fish nuclei positioned on the front.

Among the GABAergic clusters, GABA-9 and GABA-30 emerged as neural populations capable of synthesizing both acetylcholine and GABA, evidenced by their co-expression of *nkx2.1*, *lhx6* and the cholinergic markers *lhx8a*, *gbx1* and *isl1* (Fig. 2a). This molecular signature has previously been associated with neurons specifically involved in social behaviour in zebrafish ^53^. Notably, GABA-9 was uniquely characterised by the expression of genes involved in dopamine synthesis and signalling, including dopa decarboxylase (*ddc*), tyrosine hydroxylase (*th*) and dopamine-associated solute carriers (*slc18a2, slc6a3*). In contrast, GABA-30 predominantly expressed dopaminergic (*drd1a, drd1b, drd5a, thrab*) and serotonergic (*htr1b, htr1e*) receptors. This highlights a functional divergence between these two GABAergic clusters: GABA-9 appears primed for neuromodular synthesis and release, whereas GABA-30 is specialised for signal reception. A similar neuromodulatory profile to GABA-9 was also observed in the glutamatergic cluster Glut-29, which co-expressed *slc18a2, slc6a3, th* and *th2* (Fig. 2a). These transcriptomic profiles align with the known roles of dopaminergic and serotonergic signalling in regulating the motivation of cleaner fish to engage in cleaning interactions ^46,54^. Accordingly, genes involved in serotonin synthesis and signalling were co-expressed across several neuronal clusters. For instance, three GABAergic clusters (GABA-1, GABA-7 and GABA-26) expressed the dopamine transporter *slc6a3*, the tyrosine hydroxylase (*th*) enzyme involved into dopamine synthesis and various dopaminergic receptors (e.g. *drd5a*), and the immediate-early transcription factors *nr4a1* and *nr4a3*. Interestingly, these clusters also co-expressed the serotonin transporter *slc6a4*, the serotonin synthesis enzyme tryptophan hydroxylase 2 (*tph2*), and several serotonin receptors. This overlap highlights the profound cellular complexity of these neural populations in teleosts, demonstrating the possible integration of multiple neuromodulator pathways within single cell types. Among these GABAergic clusters, GABA-26 also exhibited high expression of several Immediate Early Genes (IEGs), such as *npas4a, egr1, egr2a, fosab* and *fosb,* suggesting that this cluster may represent a particularly active or highly plastic neuronal cell population. Lastly, in addition to these neuromodulatory signatures, one GABAergic cluster (GABA-14) was putatively identified as glycinergic based on the co-expression of the glycine marker genes *slc7a8* and *slc6a5*, along with the dopaminergic receptor *drd3*.

While several neuromodulatory genes were expressed across the GABAergic clusters the glutamatergic Glut-17 and Glut-2 (Fig. 2a) were also characterised by the expression of the serotonin transporter *slc6a4*, as well as serotonin receptors (*htr5aa, htr6*). Similar to the GABA-26 cluster, the glutamatergic Glut-15 cluster exhibited marked IEGs expression, highlighting the complex interplay between GABAergic inhibitory and glutamatergic excitatory signalling systems within the broader neuromodulatory network present in interacting cleaners. Lastly, the co-expression of noradrenergic-related genes (Fig. 2a), including the vesicular monoamine transporter *slc18a2* and the norepinephrine transporter *slc6a2* within the Glut-21 cell cluster, suggested a putative noradrenergic role for these cells.

### Neuroanatomical cell cluster architecture is conserved between fish species

To assign the identified cell clusters to specific neuroanatomical regions, we combined the analysis of established neuroanatomically restricted a priori genes of interest (Supplementary Excel Table S6a) with a cross-species mapping approach using SAMap, a Python-based tool specifically designed to account for protein divergence across evolutionarily distant species ^55^. Specifically, we compared cleaner fish snRNA-seq cell clusters with previously characterised cichlid snRNA-seq cell types that had been integrated with spatial transcriptomics and cross-species analyses ^56^ (Fig. 2b,c; Supplementary Excel Table S6b). Analysis of the neuroanatomical transcriptional profiles using broad regional marker genes clearly distinguished glutamatergic pallial clusters, characterised by the expression of *bhlhe22* and *eomesa,* from GABAergic subpallial clusters, which predominantly expressed *dlx2a* and *dlx5a* (Fig. S5). Among the glutamatergic pallial cells, several clusters (Glut-0, Glut-3, Glut-11, Glut-12, and Glut-16) exhibited strong expression of *pvalb7*, consistent with their assignment to the dorsolateral pallial area (Dl) ^57^. This result was further supported by the cross-species comparison, as these clusters aligned with cichlid cell clusters originating from different regions of the lateral telencephalon (Dl) (Fig. 2b). The medial zone of the dorsal pallium (Dm) has been proposed as the teleost homologue of the pallial amygdala ^57,58^. Consistent with this hypothesis, Glut-2, Glut-15, Glut-19, Glut-28 and Glut-29 displayed an *eomesa*^-^/*emx3*^+^ transcription profile ^57^ (Fig. S5), supporting their assignment to this brain region. Furthermore, SAMap analysis revealed that Glut-3, Glut-13, Glut-17, and Glut-23 strongly aligned with cichlid cell clusters assigned to the lateral telencephalon, which correspond to mouse hippocampal CA3 cell types (*bhlhe22*^+^) ^56^. These findings further support the hypothesis that teleost pallial neurons share conserved transcriptional signatures with mammalian hippocampal regions ^57,59^. Likewise, the glutamatergic cluster Glut-21 showed strong correspondence with cichlid cell clusters from the Dl area, which in cichlids exhibit transcriptional similarities to mouse neocortical cell types, such as the retro-hippocampal and cingulate/retrosplenial cortex ^56^.

The subpallial identity of the GABAergic clusters was similarly supported by their transcriptional profiles and reinforced by their SAMap alignment with corresponding cichlid cell clusters (Fig. 2b). Among these, GABA-14 and GABA-24 are putatively homologues of the mouse medial ganglionic eminence (MGE). GABA-14 co-expressed *pvalb6^+^*/*nxph1*^+^ (Fig. S5) and aligned with cichlid clusters putatively corresponding to hippocampal and cortical interneurons in mice. Meanwhile, GABA-24 expressed *sst1.1*^+^ and *npy*^+^ markers (Fig. S5) characteristic of mammalian MGE-derived inhibitory interneurons, and showed strong alignment with cichlid clusters localised to the ventral telencephalic region (Vl), supporting the assignment of these cell types to this specific brain region ^56^. In addition, GABA-1, GABA-7, and GABA-26 co-expressed *meis2*, *sp9* and *six3a* genes (Fig. S5), genes commonly enriched in mammalian striatal medium spiny neurons (MSNs) ^56^. These clusters aligned with cichlid cell clusters spatially localised to the dorsal nucleus of the ventral telencephalic area (Vd) and the supracommisural nucleus of the ventral telencephalic area (Vs), supporting the hypothesis that they represent teleost homologues of mammalian striatal MSNs. Similarly, GABA-9 and GABA-30 aligned with cichlid clusters located within the ventral nucleus of the ventral telencephalic area (Vv), a region considered homologous to the mammalian septal formation and closely related to the MGE ^37,60^. Here, the mammalian homology assignment differed from previous characterisations of the cleaner fish telencephalon (putative MGE assignment ^61^), although this may reflect the different cluster resolution used in the present study. The assignment remains putative, based on shared transcriptional profiles between cichlid clusters, their mammalian homologues and SAMap top hits. The septal assignment was further reinforced by co-expression of *isl1a, lh6a, lhx8,* and *zic1* ^56^ and by the absence of *sst1.1* and *npy*, gene markers characteristic of MGE-derived interneurons ^56^ (Fig. S5). Finally, the radial glial clusters RGc-8 and RGc-10 co-expressed canonical markers of neuronal precursors, including *tubb5*, *her4.2*, *emx2* and *emx3*, suggesting that these cell clusters contribute to the generation of pallial neurons (Fig. S5) ^57^.

### Elevated CO_2_ impact cleaner fish interaction quality

Cleaner fish and clients were exposed to either ambient or elevated CO_2_ conditions for 15 consecutive days (*n = 12* cleaners per condition) with all other conditions being the same between treatments. Analysis of the behavioural data revealed no significant effect of CO_2_ treatment on the motivation to interact across the measured parameters (Fig. 3; Supplementary Excel Table S7a,b). In contrast, regarding interaction quality, elevated CO_2_ significantly decreased the mean interaction duration (p = 0.04427; Supplementary Excel Table S7a). Post hoc comparison further revealed that this overall decrease was specifically driven by a reduction in mean cleaning duration (p = 0.0049), defined as the total time cleaners spent actively removing ectoparasites from clients, excluding periods of inspection or tactile stimulation (Fig. 3e; Supplementary Excel Table S7a,b). While we acknowledge limitations due to small sample size (*n = 12*), these results are consistent with previous studies investigating the effects of elevated CO_2_ in *L. dimidiatus* ^14^.

**Fig. 3.**
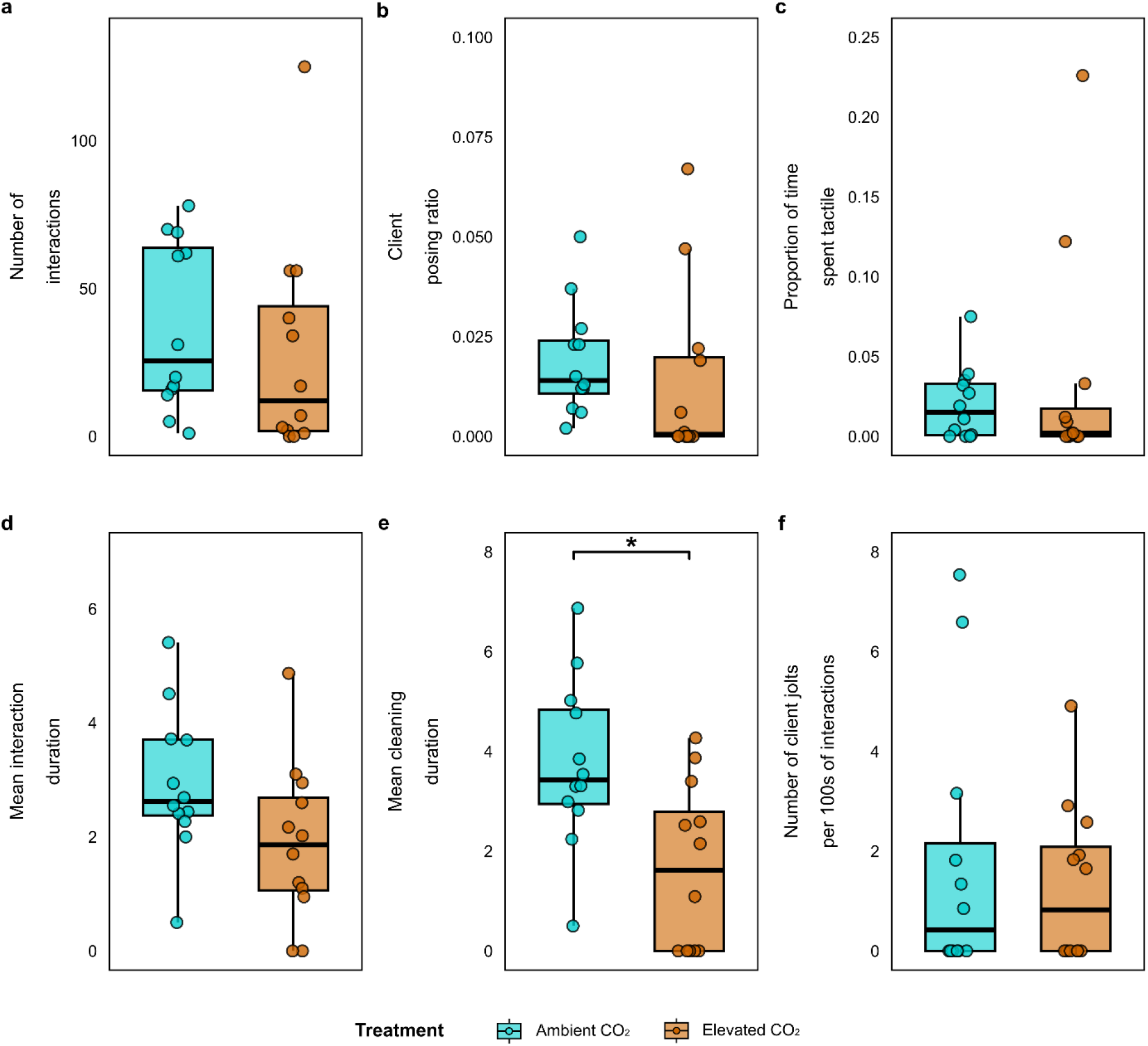
Behavioural responses elicited by elevated CO_2_. Interaction test involving cleaner fish exposed to ambient (*n = 12*) and elevated CO2 (*n= 12*). Motivation to interact is measured with **(a**) number of interactions, **(b)** client posing ratio (i.e. number of client “posing”/time not interacting). Quality of interactions are measured with **(c)** proportion of time performing tactile stimulation, **(d)** mean interaction duration, **(e)** mean cleaning duration, **(f)** number of client jolts (e.g. dishonest behaviour by the cleaners) per 100 seconds of interaction. Data presented in seconds. Treatments are represented by ambient CO_2_ (present day scenario, temperature = 25 °C, p CO_2_ ∼500 μatm), and elevated CO_2_ (near-future OA, temperature = 25 °C, p CO_2_ ∼1000 μatm). The asterisk “**\***” define statistical significance based on post hoc tests (p < 0.05). Additional details of associated behavioural analysis are found in Supplementary Excel Table S7a,b.

### Elevated CO_2_ alters GABAergic, glia and OPCs composition while inducing cell-specific transcriptional shifts

To investigate elevated CO_2_-associated changes in cell cluster composition, we analysed differences in the relative proportions of cell clusters between ambient and elevated CO_2_-treated fish. Among neuronal clusters from the ventral telencephalic area, GABA-9 and GABA-14 clusters exhibited an increased proportion of nuclei following elevated CO_2_ treatment (Fig. 4a; Supplementary Excel Table S8). Interestingly, radial glial cells, the principal source of adult neurogenesis teleost ^62^ and key mediators of behaviourally induced neurogenesis ^61,63^, also responded to elevated CO_2_ exposure, with the RGc-10 cluster showing an increased proportion of nuclei (Fig. 4a; Supplementary Excel Table S8). In contrast, OPCs, which are essential for generating myelinating oligodendrocytes and influencing neuronal structure and function ^64^, displayed a reduced proportion of nuclei in response to elevated CO_2_ (Fig. 4a; Supplementary Excel Table S8), suggesting potential vulnerabilities in neural maintenance and myelination processes.

**Fig. 4.**
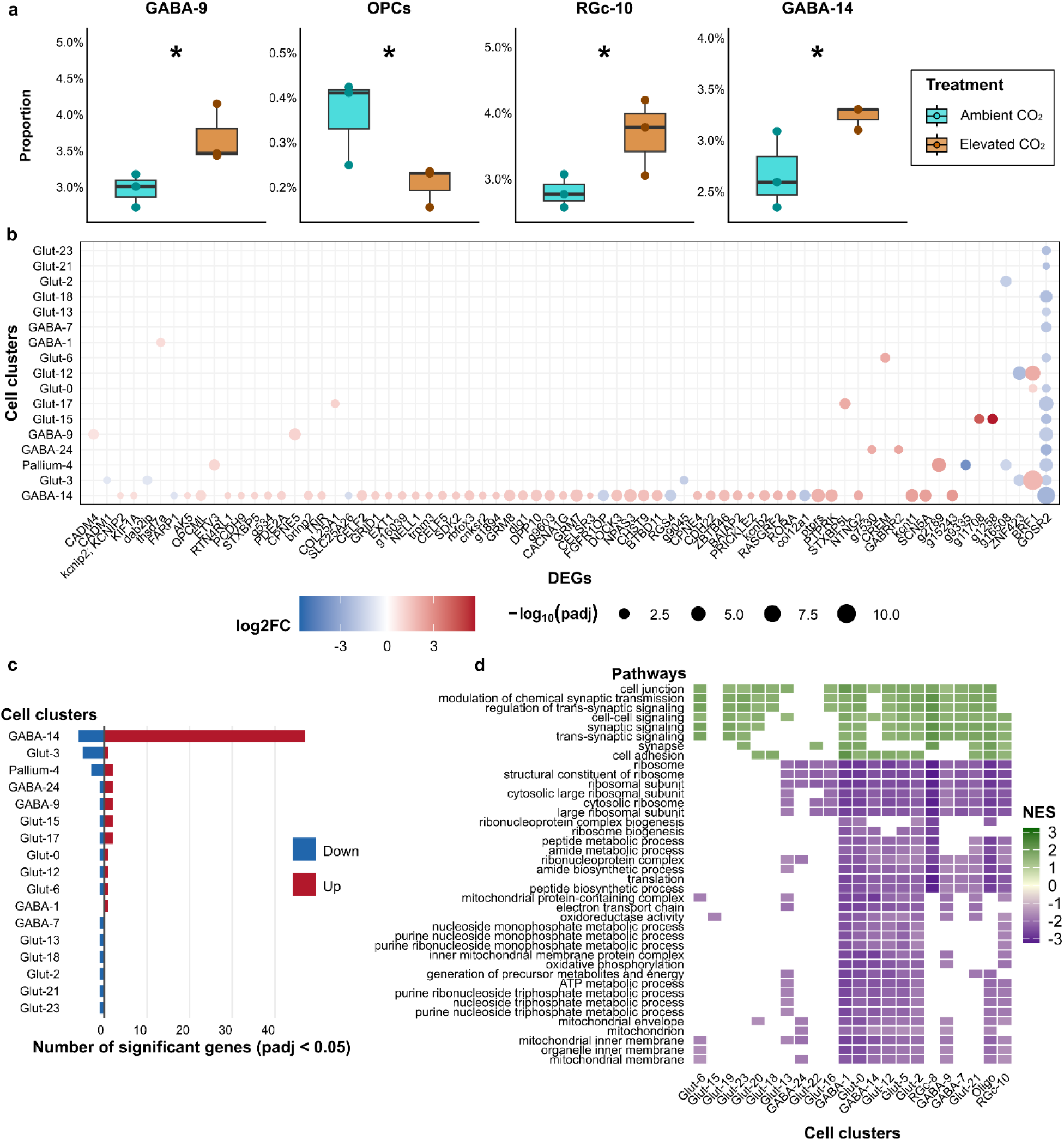
Altered cell composition and transcriptional shifts following elevated CO_2_ exposure. **(a**) Significant changes in cell cluster proportion elicited by elevated CO2 exposure (GABA-9: p = 0.0007, q = 0.0145; OPCs: p = 0.0009, q = 0.0145; RGc-10: p = 0.0016, q = 0.0173; GABA-14: p = 0.0059, q = 0.0490). See Supplementary Excel Table S8 for the results across all cell clusters. **(b)** Dotplot representing the Differentially Expressed Genes (DEGs) identified coloured based on the log2FC and sized by the negative logarithm of the adjusted p-values. Red and blue values represent upregulated and downregulated genes, respectively. For the complete DEGs list with the corresponding gene description see Supplementary Excel Table S9. **(c)** Barplot representing the number of significant DEGs (p-adjusted < 0.05) identified among all cell clusters. Red and blue values represent upregulated and downregulated genes, respectively. **(d)** Heatmap of GSEA results representing the top 40 significant pathways (padj < 0.05) enriched in distinct cell clusters following elevated CO_2_ exposure. Pathways represented are shared in equal or more than five cell clusters. Green and purple values represent high and low Normalised Enrichment Scores (NES) values, respectively with light-yellow as a midpoint; white values show pathway’s absence in the presented cell cluster. See the Supplementary Excel Table S11 for a complete list of GSEA results.

Elevated CO_2_ also induced substantial transcriptional changes in distinct neuronal cell populations. Differential expression analysis identified 89 Differentially Expressed Genes (DEGs) (FDR < 0.05) distributed across different cell-types, suggesting their potential association with the effect of CO_2_ treatment on interaction behaviour (Supplementary Excel Table S9). Among all cell types, the GABAergic cluster GABA-14 exhibited the strongest transcriptional responses with 53 DEGs, of which 47 (89%) were upregulated (Fig. 4b,c). For example, elevated CO_2_ upregulated several protein tyrosine phosphatase receptors (Supplementary Excel Table S9), a large family of transmembrane enzymes involved in regulating cell proliferation, adhesion and survival. Notably, the protein tyrosine phosphatase receptor type S (*ptprs*), which is essential for normal neural development and functions as a negative regulator of synapse formation in the adult mammalian brain ^65^ was significantly upregulated. Similarly, genes encoding for proteins critical for neurogenesis and visual system development were also upregulated, including cadherin 4, type 1, R-cadherin, protocadherin 9 (*PCDH9*), and neuronal PAS domain protein 3 (*NPAS3*) ^66–69^. Furthermore, elevated CO_2_ exposure increased the expression of genes previously associated with neurological disorders and altered excitability, such as the brain-specific angiogenesis inhibitor 1-associated protein (*BAIAP2*) and CUGBP Elav-like family member 2 (*CELF2*) ^70,71^. Given that interaction behaviour and client recognition have previously been linked to glutamatergic signalling, calcium channel activity, and immediate early genes (IEGs; Ramírez-Calero et al., 2022; von Trotha et al., 2014), the presence of DEGs across all of these functional categories within GABA-14 is particularly noteworthy. Several upregulated genes encode channel proteins with established roles in synaptic plasticity and neuronal, including the transient receptor potential cation channel subfamily M (*trpm3*) ^73^, the potassium channel subfamily T (*kcnt1*), and the potassium voltage-gated channel (*kcnb2*). Concurrently, multiple glutamate receptor genes (*GRM7, GRM8, GRID1*) were upregulated, potentially contributing to altered synaptic plasticity and cognitive function in *L. dimidiatus* ^47^. Additionally, reticulon 4 receptor-like 1a (*RTN4RL1*), an IEG-like gene recently identified in cichlids ^63^ and *L. dimidiatus* ^61^, was upregulated, further suggesting modified neuronal activity in CO_2_-treated fish. Alongside these upregulated genes, GABA-14 exhibited the downregulation of specific genes, including regulators of G-protein coupled receptor signalling and neural system development (*RGS4, FARP1*) ^74,75^, as well as collagen structural components (*col12a1*). Finally, functional enrichment analysis corroborated these transcriptional changes within the GABA-14 cluster, revealing significant enrichment for several molecular functions related to ion channel activity, calcium transmembrane transport, and glutamate receptor activity (Supplementary Excel Table S10).

Although GABA-14 exhibited the strongest molecular responses to elevated CO_2_, DEGs controlling synaptic activity and functional components were also identified in other neuronal clusters (Fig. 4b,c). For example, the GABA-9 cluster exhibited dysregulation of copine V (CPNE5) and the cell adhesion molecule *CADM4*, while GABA-24 showed upregulation in GABAergic receptor subunits (*GABRR2*). Among the glutamatergic clusters, Glut-17 displayed altered expression of collagen type XXV alpha1 (*COL25A1*) and syntaxin binding protein 5-like (*STXBP5L*). In contrast, Glut-3 showed downregulation of the cell adhesion molecule *CADM1*, the zinc finger protein *ZNF423* as well as the DAB2 interacting protein (*dab2ip*), a gene involved in multiple aspects of brain development ^76^. Lastly, it is worth noting that this analysis revealed a consistent downregulation across multiple neuronal clusters of the Golgi SNAP receptor complex member 2 (*GOSR2*) (Fig. 4b; Supplementary Excel Table S9), a key gene involved in vesicular transport from endoplasmic reticulum to Golgi ^77^, suggesting a potential specificity of the elevated CO_2_ exposure on this intracellular pathway across diverse cell types.

To further investigate coordinated transcriptional responses to elevated CO_2_ treatment, we performed Gene Set Enrichment Analysis (GSEA) on the identified DEGs. This approach revealed coordinated changes in functionally related gene sets across distinct cell types. Normalised Enrichment Scores (NES) were identified for pathways associated with synaptic functions, including synaptic signalling, transmission and organisation, and cell-cell signalling and adhesion across multiple cell clusters (Supplementary Excel Table S11). These results indicate a widespread upregulation of synaptic and neural system processes as well as ion transport activity and cell signalling in response to elevated CO_2_. Conversely, negative NES values were predominantly associated with pathways involved in ribosome biogenesis, peptide synthesis, oxidoreductase activity, ATP production as well as nucleotide/nucleoside biosynthetic and metabolic processes in response to elevated CO_2_ suggesting an extensive downregulation of the energetic and biosynthetic machinery across cell types (Fig. 4d).

### Co-expression networks reveal metabolic shifts and widespread glutamatergic overexpression

To identify the effects of elevated CO_2_ exposure on discrete cell types, we evaluated gene co-expression networks and compared their architecture between treatments using high-dimensional weighted gene co-expression network analysis (hdWGCNA) ^78^. Based on the significant changes observed in GABA-14, GABA-9, and the two identified radial glial cell clusters (RGc-8 and RGc-10) we further investigated each individually. Analysis of the co-expression networks for the two radial glial cell clusters revealed several shared pathways (Supplementary Excel Table S12a,b and Supplementary Excel Table S13a,b). These included visual learning and behaviour, exploration and social behaviour (GO:0007632, GO:0035640, GO:0035176) as well as various regulatory processes, such as the regulation of peptide and glutamate secretion, transmembrane transporter activity, axon regeneration and neurotransmitter receptor activity.

Despite these shared features, each radial glial cell cluster also displayed enrichment for distinct biological functions, indicating cell-cluster specific responses to elevated CO_2_. Among the four modules identified within the RGc-8 cluster, RGc-8-M1 and M2 were prominently enriched in terms associated with cell differentiation, migration and proliferation, and dopamine transport, respectively. In contrast, RGc-8-M3, which was positively correlated with the elevated CO_2_ treatment (Supplementary Excel Table S12c), was exclusively enriched in biological functions connected to pH regulation (GO:0006885, GO:0051453, GO:0030641), and the detection of external stimuli (e.g. GO:0009581, GO:0009593). This likely reflects a cellular response to the altered water chemistry caused by elevated CO_2_ exposure. Similarly, RGc-10-M3 which also correlated positively with elevated CO_2_ treatment (Supplementary Excel Table S13c), was uniquely enriched in terms linked to the cellular response to catecholamine stimulus (GO:0071870), cell migration and development (e.g. GO:0090497). Notably, the M3 modules from both radial glial cells were enriched in terms associated with dendrite and dendritic spine morphogenesis and development, as well as several terms related to insulin secretion and regulation (e.g. GO:0030073). They also showed enrichment for GABAergic synaptic transmission and regulation (e.g. GO:0051932) and multicellular organismal response to stress (GO:0033555). Together, these findings suggest that elevated CO_2_ exerts a direct influence on neuronal structural plasticity and stress-response pathways within these cell types.

GABA-14 exhibited the highest number of DEGs in response to elevated CO_2_ (Fig. 4b,c), with several genes indicating an increase in the excitatory state of these cells. Consistent with the pattern observed in the RGc clusters, GABA-14 showed enrichment for several terms associated with behavioural functions, including exploration behaviour, visual learning and behaviour and associative learning (GO:0035640, GO:0008542, GO:0008306), as well as organismal response to stress (GO:0033555) (Supplementary Excel Table S14a,b). This stress response was specifically enriched in GABA-14-M2, a module positively correlated with the elevated CO_2_ treatment Supplementary Excel Table S14c). In line with the increased excitatory profile suggested by the identified DEGs, GABA-14-M2, was exclusively enriched in biological functions linked to glutamate receptor signalling and transmission alongside a few terms related to GABAergic synaptic transmission, highlighting the elevated involvement of the excitatory signalling system within these cells. Additionally, this module was enriched in terms related to dendritic morphogenesis and development, insulin secretion and regulation, as well as catecholamine secretion and transport, reflecting a pattern similar to that observed in RGc-10-M3. In contrast, GABA-14-M3 and M4 were negatively correlated with the elevated CO_2_ treatment and displayed markedly distinct functional profiles (Fig. 5a,b). Specifically, M3 was exclusively enriched in functions related to the regulation of circadian rhythm and photoperiodism (e.g. GO:0042752; Fig. 5c), including core circadian clock genes (e.g. *PER2, PER3, USP2*; Supplementary Excel Table S14a), whereas M4 was enriched in terms related to sodium and neurotransmitter transport and secretion (e.g. GO:0006814, GO:0006836; Fig. 5d) suggesting that elevated CO_2_ downregulates responses related to daily oscillations and homeostasis.

**Fig. 5.**
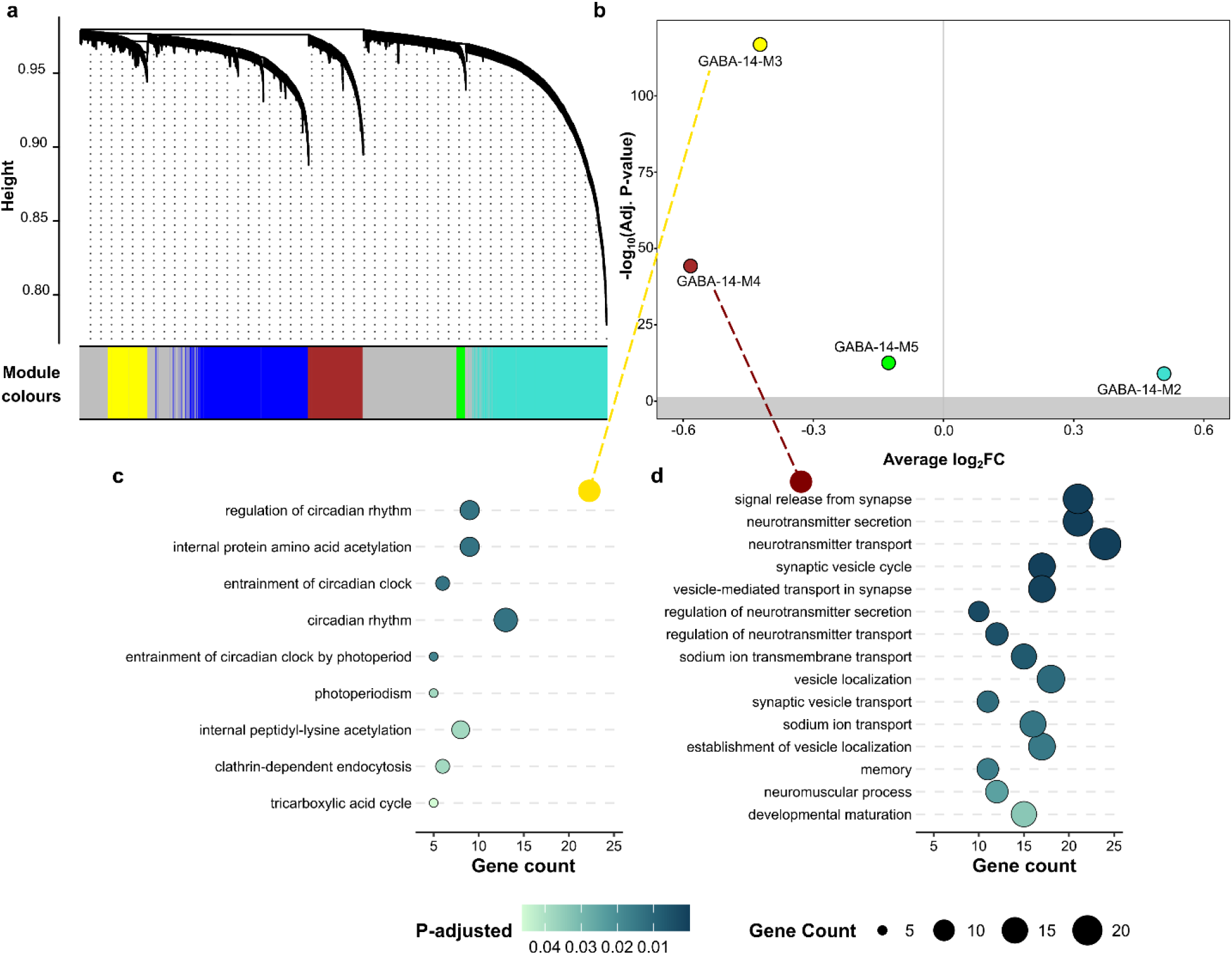
GABA-14 co-expression modules in response to elevated CO_2_. **(a)** hdWGCNA dendrogram of GABA-14 cluster. **(b)** Volcano plot of the “differential module eigengene” analysis results for the GABA-14 cluster showing the effect size (average log_2_ fold-change) and the significance (-log_10_ of the adjusted p-value). Positive and negative values are up- or down-regulated in the elevated CO_2_ exposure, respectively. Values above the grey section are considered statistically significant (padj < 0.05). (**c-d**) Top exclusive enriched biological processes ranked by adjusted p-value (q < 0.05) within GABA-14 **(c)** module 3 and **(d)** module 4. See Supplementary Excel Table S14b for a complete list of GABA-14 EnrichGO results.

Lastly, the GABA-9 and GABA-30 clusters were identified by the expression of several genes associated with social behaviour in fish (Supplementary Excel Table S15a,b). Despite the putative involvement of both clusters in social behaviour, GABA-30 was characterised by a small cell population that could introduce biases into the network analysis. Therefore, to preserve a robust cell-type-specific signal, we restricted our co-expression network analysis solely to GABA-9. Within the GABA-9 cluster, M1 and M2 modules were positively correlated with elevated CO_2_ (Supplementary Excel Table S15c), and both were enriched for pathways regulating dendrite morphogenesis and development, aligning with the pattern observed in the other CO2-associated modules. Interestingly, no terms linked to GABAergic regulation were enriched in any of the modules; instead, an extensive enrichment for glutamate receptor signalling and transmission was observed, which was particularly strong within GABA-9-M1. The M1 module was also exclusively enriched for catecholamine secretion, and insulin regulation consistent with patterns seen in other CO_2_-associated modules across the analysed cell types. Furthermore, the GABA-9-M1 module exhibited remarkably widespread enrichment for functions related to the regulation, import, and homeostasis of sodium, calcium, potassium and neurotransmitters. Together, these results suggest that elevated CO_2_ alters multiple aspects of neuronal function in GABA-9 cells, encompassing ion and neurotransmitter transport, glutamatergic signalling and neuronal structural plasticity.

### Elevated CO_2_ remodels directional cell-cell communication networks

Throughout the CellChat analysis, we identified that elevated CO_2_ induced widespread remodelling of cell-cell communication networks among neuronal cell types. Overall, both the number and the strength of cellular interactions increased in elevated CO_2_-treated fish compared to ambient controls (Fig. S6a,b). Consistent with this pattern, analysis of incoming and outgoing signalling interactions revealed an overall increase in signalling strength across all cell clusters in elevated CO_2_ (Fig. S6c,d). Given the central role of radial glial cells in adult neurogenesis in fish, their reported involvement in social behaviour ^63^, and their contribution, together with oligodendrocytes, to tissue repair following brain injury ^79^, we hypothesized that elevated CO_2_ could specifically alter communication networks between these cell types and other neuronal populations. Furthermore, because elevated CO_2_ had a pronounced effect on OPCs, which play essential roles in proliferation, differentiation, and axonal repair, we investigated communication probabilities originating from OPCs to determine whether elevated CO_2_ altered their signalling contact preferences or disrupted specific cellular interactions.

Overall, the comparison between ambient and elevated CO_2_ treatments revealed a broad reorganisation of the communication network. Although some communication probabilities increased following elevated CO_2_ exposure, others were weakened, reshaped or even completely lost (Supplementary Excel Table S16). Among the radial glial cell clusters, for example, the Sema4d-Plxnb2 signalling pathway (i.e. SEMA4 pathway) in the RGc-10 cluster established communication with the OPC cluster exclusively under elevated CO2 (Fig. 7a). The SEMA4 pathway is known for its immunoregulatory function and has been implicated in several immune-related disorders by regulating immune cell proliferation, differentiation, and migration ^80^. Conversely, although the Sema6a-Plxna4 signalling pathway (i.e. SEMA6 pathway) remained unaffected in the RGc-10 cluster (Fig. 7a), its broader communication network was substantially reorganised under elevated CO_2_. Specifically, communication with oligodendrocytes was completely lost (Fig. 6a; Fig. 7b,c), while new outgoing interactions from the RGc-8 and Glut-0 clusters emerged, alongside new incoming communication from the Glut-21 cluster. Similarly, the Lrfn4-Ptprf and Lrfn4-Ptprs signalling pathways (i.e. PTPR pathway) mediated direct communication from OPCs to multiple cell clusters exclusively under ambient CO_2_ conditions (Fig. 6b; Fig. 7d,e). The PTPR signalling pathway, which has been implicated in excitatory synapse development and locomotor behaviour in mice ^81^, was extensively remodelled following elevated CO_2_ exposure. For example, directional communication from OPCs to several glutamatergic and GABAergic cell clusters was completely abolished in the elevated CO_2_ fish, whereas signalling from the RGc-10 and Glut-23 clusters was substantially reduced but not entirely lost. Nevertheless, new outgoing interactions from both OPCs and Oligo clusters to other neuronal cell clusters involving the Nrxn3-Nlgn3 signalling pathway (i.e. NRXN pathway), a known cell communication pathway with roles in the regulation of synaptic differentiation, process information and cognitive abilities ^82^, emerged in response to elevated CO_2_ (Fig.6a,b; Supplementary Excel Table S16). Lastly, the Nrg3-Erbb4 signalling pathway (i.e. NRG pathway) exhibited an overall increase in communication probability across the analysed cell populations under elevated CO_2_ (Fig. 6a,b; Fig. 7a; Supplementary Excel Table S16). Notably, directional signalling was preferentially targeted towards GABAergic clusters previously identified as putative homologues of mouse MSNs (GABA-1, GABA-7, and GABA-26). Because the NRG signalling pathway has been associated with the regulation of dopaminergic neurotransmission, these changes may have implications for dopamine-related neural circuits and associated behaviours ^83^.

**Fig. 6.**
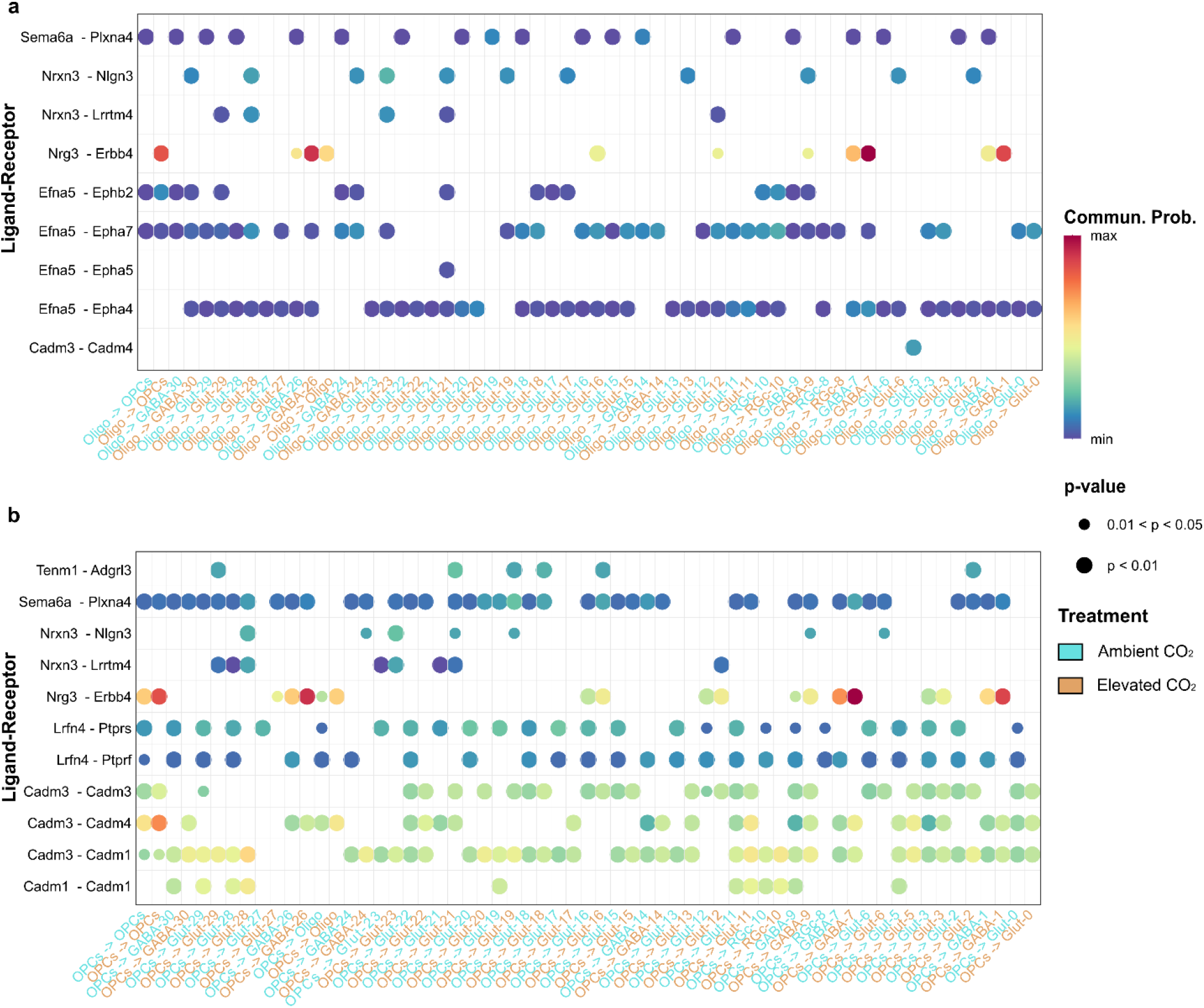
Direct cell-cell communication network probabilities are reshaped in Oligodendrocytes and Oligodendrocyte precursor cell (OPC) populations. (**a-b**) CellChat analysis comparison of ambient and elevated CO_2_ treatment conditions highlighting the directional cell communication signalling networks from (**a**) Oligodendrocytes (Oligo) and (**b**) Oligodendrocyte precursor cells (OPCs) to all other cell clusters. Only significant Ligand-Receptor pairs are displayed. Higher communication probabilities are coloured in red. For a complete list of communication probabilities and Ligand-Receptor interactions, see Supplementary Excel Table S16.

**Fig. 7.**
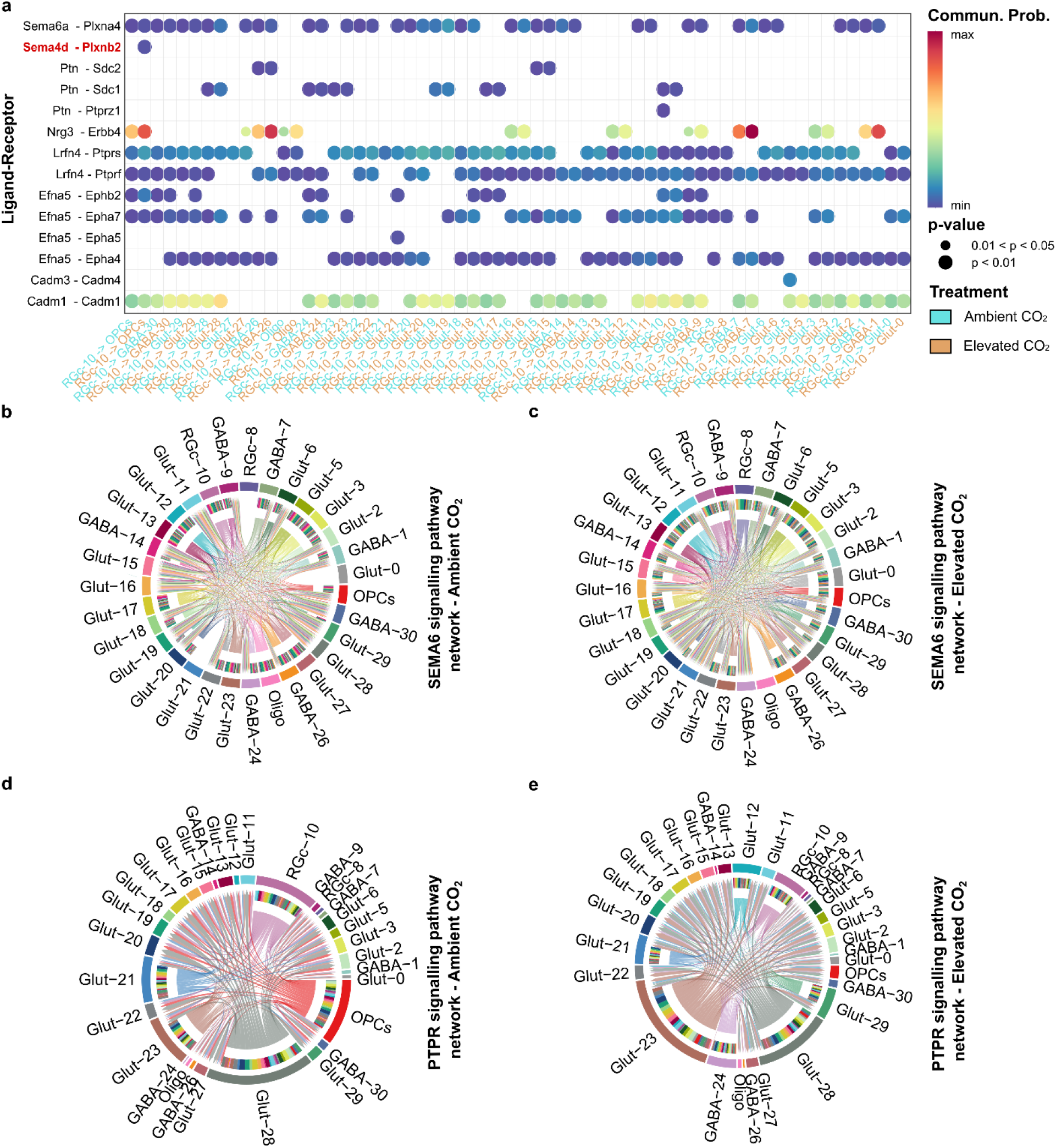
Cell-cell communication networks are reshaped in discrete cell populations. (**a**) CellChat analysis comparison of ambient and elevated CO_2_ conditions representing the directional cell communication signalling networks from radial glial cluster 10 (RGc-10) to all other cell populations. Only significant Ligand-Receptor pairs are displayed. SEMA4 signalling pathway is highlighted in bold with red colour. Higher communication probabilities are displayed in red. For a complete list of communication probabilities and Ligand-Receptor interactions, see Supplementary Excel Table S16. (**b-e**) Chord diagrams illustrating predicted cell-cell communication networks of all cell clusters involving (**b-c**) SEMA6 signalling pathway following either (**b**) ambient or (**c**) elevated CO_2_ exposure highlighting the oligodendrocytes contact lost (Oligo is coloured in pink) and (**d-e**) PTPR signalling pathway following either (**d**) ambient CO_2_ with direct communication network from OPCs to multiple cell clusters highlighted in red colour or (**e**) elevated CO_2_ exposure where this network communication from OPCs is lost.

## DISCUSSION

While elevated CO_2_ is known to alter both brain gene expression and mutualistic behaviours in marine fishes ^14,35^, the precise cellular target underlying these cognitive and behavioural shifts have remained unresolved. Here, we provide the first high-resolution cellular atlas of the telencephalon in the highly social interacting cleaner wrasse *L. dimidiatus* under projected near-future elevated CO_2_ conditions. By simultaneously profiling heterogeneous cell populations we reveal that environmental acidification does not merely induce a uniform neural stress response; rather, it triggers highly cell-specific vulnerabilities. Specifically, we show that elevated CO_2_ alters glial population dynamics, marked by depletion of OPCs, and elicits transcriptional reprogramming in social and circadian associated neuronal clusters. Ultimately, these targeted cellular impacts culminate in a widespread disconnection and remodelling of directional cell-cell communication networks, providing a mechanistic cellular basis for the altered mutualistic interactions observed in acidified oceans.

As a critical foundation for our molecular findings, we first confirm that elevated CO_2_ exposure affected the duration of cleaning interactions in our behavioural tests. Consistent with previous studies ^14,35^, the treatment significantly reduced the time cleaner spent actively removing ectoparasites from client fishes. Since cleaners are known to influence local fish communities ^26^, such individual behavioural changes could have broader ecological consequences, potentially disrupting mutualistic relationship with clients and, ultimately, affecting reef community dynamics. Anchoring these macro-level behavioural alterations to precise neural circuits requires a robust neuroanatomical framework. Our classification of the cleaner fish telencephalon provides such a framework, revealing a deeply conserved cellular architecture broadly consistent with findings from cichlids ^56,63^, zebrafish (*Danio rerio*) ^57^ and goldfish (*Carassius auratus*) ^59^, as well as with recent transcriptomic data from this fish species ^61^. By successfully assigning these discrete pallial and subpallial cell clusters to their putative mammalian homologues, we established the high-resolution map required to pinpoint the specific cell-targeted consequences of elevated CO_2_ exposure.

With this high-resolution map, Oligodendrocyte Precursor Cells (OPCs) emerged as a notable vulnerable population, exhibiting marked decrease in their relative nuclei proportion under elevated CO_2_ conditions. OPCs are highly dynamic cells that generate new oligodendrocytes throughout the lifespan, synthesising myelin sheaths required for proper axonal and synaptic function ^64^. Beyond myelination, they actively regulate axonal and synaptic development, participate in immune signalling and migrate to containment of axonal injury ^64^. Besides these functions, OPCs ablation in zebrafish optic tectum disrupts visual processing behaviour resulting in reduced prey-capture success ^84^. Because cleaner fish rely on visual discrimination and learning abilities to recognise clients and maintain mutualistic relationships ^36^, the elevated CO_2_-induced depletion of OPCs could have profound neuroethological consequences. A reduction in myelinated oligodendrocytes would likely disrupt the neural circuitry underlying visual processing, thereby compromising the cleaners’ ability to identify clients and engage in social interactions.

Amplifying this potential structural loss, elevated CO_2_ caused a functional rewiring of OPC-mediated network signalling. Specifically, we observed the *de novo* emergence of directional cell-cell communication from OPCs to several neuronal clusters exclusively in elevated CO_2_-exposed fish. This novel signalling included the NRXN pathway, involving the synaptic cell-adhesion molecule Nlgn3, which is an important regulator of cognitive behaviour. Its dysregulation can compromise the delicate balance between excitatory (glutamatergic) and inhibitory (GABAergic) signalling ^82,85,86^ and induce fear conditioning responses ^87^. On the contrary, the PTPR pathway, which is critical for neural circuit assembly, was completely abolished in cleaners exposed to elevated CO2. The inferred PTPR signalling suggests a substantial disruption of locomotory and excitatory networks, consistent with its role in regulating excitatory synapse and axon development in mammals ^81^. Although direct experimental evidence linking OPC depletion, structural myelin loss, and excitatory-inhibitory imbalance to behavioural shifts in cleaner fish requires further exploration, our data strongly suggest that OPCs reduction and the resulting alterations in neuronal signal transmission represent primary cellular responses driving the effects of ocean acidification on mutualistic interaction behaviour.

In contrast to OPCs, radial glial cells (RGc-10), alongside specific GABAergic populations (GABA-9 and GABA-14), exhibited an increased relative nuclei proportion following elevated CO_2_ exposure. Radial glia (RG) are central drivers of adult neurogenesis in teleosts ^88^, and play roles in mediating neuronal production and circuit remodelling in response to complex social cues as seen in in cichlids ^63^ and interacting cleaner fish ^61^. Adult zebrafish RG rapidly proliferate in response to neuronal loss, and their dysfunction has been associated with neurodevelopmental disorders ^62^. Hence, the expansion of RGc-10 under elevated CO_2_ could represent an initial compensatory, neurogenic response aiming at mitigating neuronal damage and maintaining telencephalon cellular homeostasis. However, our cell-cell communication analysis reveals that this glial proliferation likely comes at a cost to OPC survival. Under elevated CO_2_, the RGc-10 cluster exclusively activated the SEMA4D pathway, uniquely targeting the OPC cluster. SEMA4D is a crucial driver of neuroinflammation that can compromise blood-brain barrier integrity ^80,89^. Notably, SEMA4D is also know to directly inhibit OPC migration and differentiation ^89^. The *de novo* emergence of this RG-to-OPC SEMA4D signalling provides a compelling mechanistic link between our observations: the CO_2_-induced potentially inflammatory expansion of radial glia appears to actively supress OPC maturation, thereby intensifying the loss of mature, myelinating oligodendrocytes necessary neural circuitry underlying normal social processing.

Returning to the neuronal populations affected by elevated CO_2_, the expansion of the GABA-9 cluster is particularly noteworthy. This cholinergic-GABAergic cell population co-expresses *nkx2.1*, *lhx6* and the cholinergic markers *lhx8a*, *gbx1* and *isl1*, a transcriptional profile resembles that of a zebrafish (*Danio rerio*) telencephalic population that governs social orientating behaviour ^53^. This homology implicates GABA-9 in the regulation of cleaner fish social interactions. Interestingly, while a related cluster (GABA-30) shared this broad transcriptional signature, its proportion remained unaffected by elevated CO2. The critical distinction between the two populations lies in their neuromodulatory profile: whereas GABA-30 is specialised for signal reception, GABA-9 is distinctly primed for dopamine synthesis and release.

Dopamine is a well-established driver of motivation in cleaner fish mutualisms ^46,54^. Because previous studies have demonstrated that elevated CO_2_ reduced dopamine concentrations within discrete brain areas of cleaner fish ^11^, the observed expansion of GABA-9 could reflect a compensatory mechanism. By scaling up the cellular capacity for dopamine synthesis the telencephalon may be attempting to preserve dopaminergic homeostasis and sustain cleaning motivation under elevated CO_2_ conditions. This hypothesis is further supported by the expression of *slc18a2* (VMAT2) within GABA-9. While typically a classical vesicular transporter for dopamine ^50^, *slc18a2* can also co-package and release GABA, indicating an interplay between GABAergic and dopaminergic co-transmission. Ultimately, this delicate neurochemical balancing act is vital for maintaining the reward processing, visual stimulus evaluation and risk perception required for successful mutualistic interactions, potentially explaining why the behaviour is altered rather than completely abolished under elevated CO_2_.

Beyond shifts in cellular proportion, the GABA-14 cluster underwent profound transcriptional reprogramming under elevated CO2. Previous bulk-tissue transcriptomics demonstrated that ocean acidification alters forebrain networks in cleaner fish, specifically modifying glutamatergic signalling, calcium channel activity, visual perception, and immediate early gene responses ^14^. Our single-cell atlas resolves broad transcriptomic shifts to a more precise cellular category: remarkably, these functional pathways were concentrated and dysregulated within GABA-14 cluster. The upregulation of genes driving neuron system processes and glutamate receptor signalling points toward an alteration of synaptic plasticity. This functional inference is reinforced by our cross-species mapping which aligns GABA-14 with mammalian hippocampal and cortical interneurons, cell populations fundamental for normal brain functions and memory processes ^90^. Because cleaner fish rely heavily on sophisticated memory and learning abilities to recognise clients ^36^, the transcriptional remodelling within this specific cell population likely represents a core neural substrate for the cognitive disruptions induced by elevated CO_2_.

Beyond their established roles in learning and memory, hippocampal and cortical interneurons are integral to the regulation of intrinsic circadian rhythms ^91^. This dual function provides an intriguing physiological link to another major consequence of elevated CO_2_ in the GABA-14 cluster: the alteration of visual processing. Our co-expression analysis isolated this effect to a specific gene module (GABA-14-M3), which was negatively correlated under elevated CO_2_. This module is anchored by core circadian clock hub genes including *PER2*, *PER3* and *USP2*. At the molecular level, *USP2* modulates central circadian rhythmicity by interacting with other clock proteins such as *PER2* ^92^, which itself acts as a central regulator of visual behaviour and visual information processing ^93^. Alongside the alteration of these clock genes, markers crucial for neural retina survival and differentiation, such as cadherin 4 type 1 (also known as R-cadherin) ^66,69^, were dysregulated within GABA-14. Consequently, terms related to circadian clock entrainment were uniquely enriched within this module, suggesting a coordinated alteration in the cleaners’ ability to process external cues. Circadian rhythms and visual inputs are constantly integrated to regulate and synchronise animal behaviours ^94^. While previous studies have observed broad CO_2_-induced circadian and visual alterations across several teleost species ^8,16,95^, our findings specifically map this susceptibility to GABA-14 interneuron population. Although causal relationships underlying these mechanisms needs to be determined, because cleaner fish rely on precise visual perception and temporal regulation to execute ecologically vital behaviours, such as feeding, predator avoidance, and client recognition, the impairments of these specific networks likely play a direct role in driving the reduced cleaning duration observed under ocean acidification.

Living in an environment characterised by elevated CO_2_ forces fish to continuously adjust their internal pH homeostasis, an inherently energy-demanding process that relies on active ion transport ^8,96^. Our co-expression analysis captured the molecular footprint behind this metabolic demand, emphasizing the positive correlation of elevated CO_2_ with pathways governing insulin secretion and regulation across multiple cell clusters, potentially reflecting the increased metabolic cost required to maintain pH homeostasis. For example, module 3 of the radial glia cluster (RGc-8-M3) was positively correlated with pH regulation, while GABA-9 module 1 (GABA-9-M1) prioritised the transport and homeostasis of ions and neurotransmitters. To support these critical homeostatic processes, the brain appears to sustain an energetic trade-off. Gene set enrichment analysis (GSEA) revealed a systematic downregulation of functionally foundational pathways, including ribosome biogenesis, peptide synthesis and ATP production. This biosynthesis depression was accompanied by a pronounced and widespread downregulation of *GOSR2*, a key mediator of vesicular trafficking between the endoplasmic reticulum (ER) and the Golgi apparatus ^77^. Because environmental stressors are known to trigger ER stress in cleaner fish ^14^, this widespread suppression possibly impairs intracellular protein-processing, further compromising cellular homeostasis to redirect resources towards maintaining essential acid-base balance.

Despite these apparent energetic constrains, elevated CO_2_ surprisingly triggered a substantial upregulation in neuronal excitatory signalling. GSEA identified widespread enhancements in synaptic signalling, whilst co-expression modules characterised by glutamate receptor transmission, and dendritic spine morphogenesis where positively correlated with the elevated CO_2_ treatment. While dendritic spines are key structural components regulating synaptic plasticity, which in turn can contribute to learning and memory ^97^, their development and dynamics are also highly sensitive to stress-related factors, alongside a close interconnection with nervous and immune system activation ^98^. Indeed, the broad upregulation of pathways related to organismal stress response and catecholamine secretion suggests that even after two weeks under elevated CO_2_ exposure, the fish remain in a state of sustained physiological stress. Persistent activation of these pathways could contribute to aberrant alterations in neuronal circuitry, a hypothesis consistent with observations in other teleost where elevated CO_2_ induced anxiety-like behaviours remain altered even after prolonged recovery periods ^7^. Taken together, the concurrent evidence, raised ion-regulatory demands and suppressed biosynthesis, along with enhanced excitatory signalling and persistent stress responses, illustrate the physiological pressure imposed by ocean acidification. To maintain basic physiological equilibrium, cleaner fish need to redirect cellular resources towards homeostatic maintenance, a systematic metabolic and neurological rewiring that ultimately compromises their behavioural regulation and mutualistic interactions.

Collectively, our findings demonstrate that CO_2_-induced behavioural alterations do not emerge from the localised failure of a single neurotransmitter system, but rather from systemic, coordinated shift across diverse neural circuits. By resolving the brain’s response at single-cell resolution, we captured distinct transcriptional shifts and disconnected cell-cell communication networks that can compromise sensory integration and social interaction strategies. Notably, non-neuronal populations, such as OPCs emerged as critical vulnerabilities under elevated CO_2_, likely driving cascading effects across broader signalling pathways. Ultimately, this study provides a reproducible molecular signature linking alteration in specific cell types and communication networks to the modification of complex, ecologically vital mutualistic behaviours. By providing a foundational single-cell perspective on how ocean acidification rewires the teleost brain, these results establish a critical framework to determine whether these cellular shifts represent transient physiological coping mechanisms or permanent, maladaptive alterations to neural circuitry.

## MATERIALS AND METHODS

### Fish husbandry

Adult cleaner wrasses *Labroides dimidiatus* and client surgeon fish *Zebrasoma scopas* ^99^, were obtained from a local fish distributor (Sealife Hong Kong Limited) and delivered to The Swire Institute of Marine Science (SWIMS) aquarium. Each cleaner fish was kept in separate individual tanks (30 L) to avoid aggression among individuals, whereas one to three clients were kept together in each 50 L aquarium. Cleaner fish were fed *ad libitum* once per day with mysis frozen shrimps (Hikari Bio-Pure®) and the spirulina (Tropical®) was used for clients. Similarly to other studies on the same species ^14^, fish were acclimated for three days to the aquarium conditions before the start of any treatment. Following acclimation, in line with previous studies on the same species ^35^, CO_2_ was gradually increased over one day, and each fish was then exposed for 15 consecutive days to (1) present day scenario (ambient CO_2_, control treatment; cleaners *n = 12*; clients *n = 6*; salinity at 35 ppt, temperature 25°C, pH 8.0 ± 0.06 and pCO_2_ ∼ 500 μatm), or (2) elevated CO_2_ (continuous elevated CO_2_ treatment; cleaners *n = 12*; clients *n = 6*; salinity at 35 ppt, 25°C, pH 7.8 ± 0.03 and pCO_2_ ∼ 1000 μatm) following the 2100 projections of the IPCC’s RCP scenario 8.5 (“IPCC,” 2023). Because behavioural impairments following elevated CO_2_ exposure have been reported after only four days, the exposure duration selected in this study was consistent with previous studies in cleaner fish as well as other fish species and sufficient to elicit behavioural responses ^35,41,42^. Individual tanks had a semi-closed flow-through aquatic system where alkalinity levels, dissolved carbons and pH were monitored daily and kept constant. OA-like conditions were automatically adjusted by direct injection of CO_2_ gas (Harvest Wise Gases Hong Kong Limited; purity 99.5%) previously mixed with atmospheric air inside a mixing tank. Tanks were kept under a photoperiod of 12h/12h (light/dark cycle) with water temperature kept constant. Ammonia and nitrate levels were checked twice a week using colorimetric tests (Sera). To complement the automatic system, an additional manual daily monitoring of seawater temperature, dissolved oxygen (Pro 2030), salinity (portable refractometer), and pH (pH meter, BWA-2018SD) was used. Finally, total alkalinity (TA) was measured every other day by equivalent point titration using a compact titrator G20 (Mettler Toledo) with a TA batch 202 as reference for oceanic CO_2_ measurements (Scripps Institution of Oceanography). Using the titrator’s measurements, seawater carbonate chemistry parameters were calculated in CO2SYS v2.1 (https://www.ncei.noaa.gov/access/ocean-carbon-acidification-data-system/oceans/CO2SYS/co2rprt.html) from TA and pH measurements, similarly reported on previous studies ^15,101^. Seawater carbonate chemistry, tank parameters, and information on fishes can be found in the Supplementary Excel Table S17a,b.

### Behavioural experiment

Behavioural tests were performed in independent observation tanks (50 x 30 cm and 25 cm high; 50 L) in an isolated space. On the day of the experiment, cleaner fish were placed into an observational tank with either (1) interaction in present day conditions (cleaner-client pair in ambient CO_2_ water) or (2) interaction in elevated CO_2_ conditions (cleaner-client pair exposed to elevated CO_2_) (Fig. 1a). Before the start of the behavioural test, fish were first acclimatized to the observation tank for 20 minutes with an opaque partition between the two fish. Then, the partition was raised, and behaviour was recorded by a video camera (GoPro Hero 10 black) positioned on the front of the tank for 18 minutes, in agreement to previous studies in this species ^102^. Clients were similar in size and colour pattern for both ambient and CO_2_ conditions (according to the fish supplier catalogue were 6-10 cm size) and they interacted with one cleaner fish per day only. To avoid biases in gene expression related to food metabolism cleaners underwent a 24 h fasting period before the interaction test ^14^.

### Statistical analysis of behavioural data

Behavioural videos were blindly inspected as reported in previous studies ^11,14,47^ investigating both motivation to interact and interaction quality of cleaners. Briefly, the number and proportion of interaction started by cleaners (i.e. close body inspection and cleaning activity to remove damaged tissue or scales), and the client “posing” ratio performed to attract cleaners (i.e. number of client “posing”/time not interacting) were measured as indicative of motivation to interact. While the duration of interaction events, the proportion of time performing tactile stimulation (i.e. cleaners using their pectoral fins to “massage” the clients to reduce the stress and prolongate the interactions), and the frequency of client jolts per 100s of interaction (representative of a dishonest behaviour by the cleaners) were measured as indication of interaction quality.

For the video analysis, the event logging software “Boris” v8.27.10 ^103^ was used considering both cleaner and client fish as focal subjects (behavioural catalogue and behavioural results detailed in Supplementary Excel Table S18a,b). A total of 24 individuals were included for the statistical analysis (*n = 12* interaction in elevated CO_2_; *n = 12* interaction in ambient CO_2_ water) and data exploration was performed as in Zuur et al., 2010. Behavioural data were analysed using Generalised Linear Models (GLM) setting the treatment (factor with two levels: ambient control vs elevated CO_2_) and batch effect (factor with two levels: batch 1 and batch 2) as categorical fixed factors. In these models, Gaussian distribution was used for continuous data (interaction duration, client posing ratio and jolts ratio), the negative bimodal distribution for count data (number of interactions, since overdispersion of Pearson residuals was observed using Poisson distribution) and binomial distribution for proportions (proportion of interactions with tactile stimulation). Model assumptions (independence and absence of residual patterns), were verified by plotting residuals against fitted values and each covariate in the model. The full model was tested using the function “lme4” and the Anova Type II Wald chi-square tests from the package “car” ^105^ in R v4.4.2 ^106^. Post hoc multiple comparisons were performed using the package “emmeans” ^107^ with Tukey corrections.

### Tissue dissection and nuclei isolation

Following the behavioural test, cleaners were immediately euthanized by severing the spine, measured for standard length and body weight, and brain were dissected from the skull under a stereomicroscope (Supplementary Excel Table S17b). Telencephalon areas were dissected in ice cold Neurobasal medium (with 2% B27 and 0.5 mM Glutamax; Thermo Fisher Scientific) containing 0.2 U/μl RNase Inhibitor (Sigma), rapidly frozen in liquid nitrogen and stored at -80°C until further process. All procedures were following the ARRIVE guidelines and were approved by the Committee on the Use of Live Animals in Teaching and Research (CULATR) of the University of Hong Kong and by the Department of Health of the Hong Kong Government (Protocol and Ref. numbers: #22-051, #24-676).

Three samples per treatment were randomly selected and further processed to isolate the nuclei (*n = 3* interaction in ambient CO_2_ water*; n = 3* interaction in elevated CO_2_) by using a protocol adapted from the 10x Genomics Demonstrated Protocol CG000393 • Rev A (https://cdn.10xgenomics.com/image/upload/v1660261285/support-documents/CG000393_Demonstrated_Protocol_Adult_Mouse_Nuclei_Isolation_RevA.pdf) and fully optimized for *L. dimidiatus* telencephalon. Briefly, each frozen telencephalon was treated with lysis buffer containing 10 mM UltraPure Tris-HCl, pH 7.5 (Invitrogen), 10 mM NaCl (Invitrogen), 3 mM MgCl_2_ (Invitrogen), 0.1% NP-40 Surfact-Amps™ Detergent Solution (Thermo Scientific), 1% BSA (Sigma), 0.2 u/ul of Protector RNAse inhibitor (Roche) and RNase-free water and incubated on ice for 2 minutes. After the incubation, a sterile pestle was used to finely homogenise the tissue and the obtained lysate was completely dissociated using a P1000 pipette and left on ice for other 10 minutes. To remove large debris and nuclei aggregates formed during the dissociation step, the lysate was transferred throughout a 40 μm filter (Genever®-CS04) and centrifuged (600 rcf; 5 min; 4°C). The resulting supernatant was discarded, and the pellet was resuspended in 1.5 ml chilled wash and resuspension buffer (dPBS (Gibco) with 2% BSA (Sigma), 0.2 u/ul of Protector RNAse inhibitor (Roche)). An additional filtering step with a 10 μm filter (pluriSelect) was applied to the nuclei suspension to remove small debris and nuclei aggregates and then centrifuged (500 rcf; 5 min; 4°C). Purified nuclei were finally resuspended in 150 μl of wash and resuspension buffer to reach a final concentration between 700-1900 nuclei/μl, counted in a disposable haemocyte (Millicell®) using the Ethidium Homodimer-1 staining dye (Thermo) at the fluorescent microscope (Nikon DS-Ri2 camera). Nuclei were then immediately processed to the 10x Genomics platform.

### Single-nuclei RNA sequencing and quality pre-processing

The purified nuclei suspensions were processed with 10x Genomics Chromium Next GEM Single Cell 3ʹ Reagent Kits v3.1 (Dual Index; User guide GC000315, Rev F) and Chromium Next GEM Chip G Single Cell Kit with a nuclei target recovery of 10,000 per sample. Single nuclei were encapsulated into Gel-Beads-in-emulsion (GEM) by 10x Chromium iX according to the manufacturer manual guidelines. Library were constructed and sequenced on an Illumina NovaSeq 6000 (paired ends, 150 bp) with a total throughput of ∼100Gb at the Centre for PanorOmic Science (CPSO), the University of Hong Kong.

Raw FASTQ files were subsequently analysed with 10x Genomics Cell Ranger v6.0.1 pipeline ^108^, and the splice aware alignment tool (STAR) was used to align the reads to the *Labroides dimidiatus* genome assembly (ASM3071049v1) within Cell Ranger using the flag *--include-introns* on. Gene and functional annotations for *L. dimidiatus* were custom obtained from the same genome assembly (see Desantis *et al.* ^61^, for a complete genome annotation pipeline). The final annotation file used in this study can be found at “https://www.schunterlab.com/resources”. As a standard filtration procedure, UMIs containing homopolymers, N values, or with a base quality score less than 10 were filtered out by CellRanger. Overall, we sequenced 75,248 nuclei at an average depth of 34,724 reads/nuclei with a percentage of reads mapped confidently to both the genome and transcriptome ranging between 84.1-85.5% and 34.6-44.2%, respectively (Supplementary Excel Table S1a).

Data were imported into R v4.5.2 and each library was quality inspected independently. SoupX v1.6.2 ^109^ programme was used with default settings to remove ambient RNA. Low quality cells were filtered with the following settings: nFeatures above 250 and below 5000; % of mitochondrial genes < 3, and doublets were removed using scDblFinder v1.24.10 ^110^ using default parameters. Quality control metrics for each library are provided in the Supplementary Material (Supplementary Excel Table S1b). In total, 66,878 cells passed the quality filters and were processed for downstream analyses.

### Normalization, data integration and cell cluster identification

For data normalization, a standard SCTransform v2 pipeline within Seurat v5.4.0 ^111^ was used with a Gamma-Poisson Generalized Linear Model and by regressing out the percentage of mitochondrial reads ^112^. Each individual sample was linearly reduced using “RunPCA” function with 50 principal components (PCs). Based on the Elbow plot inspection, 30 PCs were included for the Uniform Manifold Approximation and Projection (UMAP) “RunUMAP” and “FindNeighbours” functions.

The “IntegrateLayers” function with the reciprocal PCA analysis (RPCA) was used for data integration and clusters were computed on the integrated object by employing the “FindCluster” function (*res = 1.3*). To determine the appropriate resolution, Clustree v0.5.1 package ^113^ was used to plot the relationship between clusters at different resolutions. To provide a biological identity of the identified cluster, we used both unbiased analysis of cluster specific marker genes with the “FindAllMarker” function (*only.pos = TRUE, min.pct = 0.25, logfc.threshold = 0.25, test.use = “wilcox”*) and well-established cell type specific marker genes from literature (Supplementary Excel Table S3) ^56,57,59,62,63,114^ as well as marker genes from CellMarker 2.0 ^115^. Lastly, to provide a comprehensive investigation of the transcriptional relationship of the identified clusters, we performed a cell type tree reconstruction based on pairwise expression distances between cell clusters using 10,000 bootstraps and considering genes expressed only if the average expression was greater than three transcript per million in at least one identified cluster ^116^.

### Assignment of ortholog genes

To identify the ortholog genes between *L. dimidiatus*, zebrafish and mouse species, we used diamond v2.1.24 ^119^ and we performed BLAST analysis. The protein sequences of zebrafish (*D. rerio*; genome assembly used: GCF_049306965.1) and mouse (*M. musculus*; genome assembly: GCF_000001635.27) were used with the following parameters *-f 6, --sensitive*. Only the gene with the highest percent identity match was retained if there were one-to-many or may-to-one matches. Zebrafish gene symbol annotation is reported throughout the manuscript and within all figures to achieve consistency and easier interpretation.

### Neuroanatomical inference and intraspecific single nuclei comparison

To provide insights into the neuroanatomical origins of the cleaner fish cell clusters, the expression profile of known neuroanatomically restricted a priori genes of interest (e.g. pallial and subpallial marker genes; Supplementary Excel Table S6a for a complete list) was examined. As a second line of investigation, the SAMap python tool ^55^ was used to perform a cross-species comparison of distantly related species. Here, the cichlid *Mchenga conophoros* ^63^ snRNA-seq data was selected as a teleost reference to compare transcriptional similarities of cell types by utilising the availability of both single nuclei and spatial transcriptomic data, which allowed a deeper comparative investigations. Briefly, the protein sequence from the same reference genome assembly used to create the cichlid snRNA-seq dataset (GCF_000238955.4) was downloaded and the custom SAMap’s map_genes.sh bash script was used to identify reciprocal blast hits between cleaner and cichlids protein sequences. The raw gene count matrix from cleaner and cichlid fish were converted into the accepted SAMap’s h5ad file format and the data were finally processed with the “run” function from SAMap by specifying the gene symbols of each protein for each fish species and using default parameters. The GenePairFinder function from SAMap was used to further determine the genes driving cell type relationships with default settings. We retrieved all genes considered to positively contributing to the cross-species correlation of cell types (Table S19). All cross-species mutual top hit cell type pairs have been considered as homologous. The neuroanatomical locations was inferred based on the SAMap results from this study as well as the spatial mapping of the cichlid cell clusters and the cross-species comparison reported in a the published study from Hegarty et al., (2024).

### Cluster proportion analysis

The cell proportion between clusters was compared between ambient and near-future OA treatment using a mixed effect binomial regression model employing the “glmer” package in R. Within the model, treatment conditions were a fixed effect, fish individuals were set as random effect and nuclei from either the cluster of interest or not were considered as the outcome variable. P-values were adjusted with the Benjamini-Hochberg correction Differences (significance at q < 0.05).

### Differential gene expression analysis

To identify differentially expressed genes elicited by the CO_2_ treatment, we used DESeq2 v1.50.2 ^120^ with a pseudobulk approach which treat samples as independent observations to avoid pseudo replication issues. Gene counts of all cells from the same sample for each cell type were summed by using the “AggregateExpression()” function from Seurat ^111^. Lowly expressed genes (present in less than 5% of cells) were filtered out. The Wald statistic test was used considering the treatment (CO_2_ vs Int) within the formula and identified genes with a False Discovery Rate (FDR) p-adjusted values of < 0.05 were considered differentially expressed.

### Gene ontology enrichment analysis

To identify the major functional processes within the identified cell cluster, functional enrichment analysis of Gene Ontology (GO) categories among cluster-specific differentially expressed genes (DEGs) was investigated. The AnnotationForge v3.22 ^117^ package was used to build the genome-wide annotation (OrgDb) containing all specific GO for *L. dimidiatus*. The GO pathways were retrieved from the *L. dimidiatus* annotation file (https://www.schunterlab.com/resources). Then, the profiles of each gene cluster were computed using the “enrichGO” function provided by ClusterProfiler v4.18.4 ^118^ and FDR-adjustment (*q* < 0.05) were considered statistically significant.

### Gene set enrichment analysis (GSEA)

The unsupervised GSEA with the fgsea R package v1.37.4 was used on the DEGs results from DESeq2. A ranked gene list containing all the DEGs identified was supplied and GO pathways were retrieved from the *L. dimidiatus* annotation file (https://www.schunterlab.com/resources). Normalised enrichment score (NES) values were calculated, and functions were considered significant with p-adjusted < 0.05.

### Cluster specific high-dimensional weighted correlation network analysis (hdWGCNA)

The hdWGCNA analysis ^78^ was performed using the neuronal cell clusters GABA-9, GABA-14, and the radial glial cell (RGc-8 and RGc-10) clusters separately to identify gene modules associated with the CO_2_ treatment. For each of the analysed clusters, soft thresholding values were selected using the “TestSoftPowers” function (GABA-9 and GABA-14: *6*; RGc-8: *5*; RGc-10: *4*) and the co-expression network was built with the “ConstructNetwork” function. For each module, module eigengenes and module connectivity were computed using default settings and gene expression within each module was identified. Genes were ranked by intramodular connectivity (kME) to identify highly connected hub genes within each module. As the focus of the analysis was the identification of molecular pathways connected to the CO_2_ treatment and neural signalling, we excluded from further analysis modules (RGc-10-M4; GABA-9-M3; GABA-14-M1) showing hub genes primarily enriched in mitochondria activity (Supplementary Excel Table S13a, 14a, 15a). The differential module eigengene (DME) analysis was used to discover modules with significant positive and/or negative correlations with the treatment (*min.pct = 0.25*). ClusterProfiler v4.18.4 ^118^ with the “enrichGO” function was used to perform the overrepresentation analysis and results were considered significant with FDR-adjustment of *q* < 0.05. To reduce the functional redundancy, the web programme “REVIGO” ^121^ was used with default parameters to inspect the overrepresented pathways results.

### Inference cell-cell interactions

To identify major signalling changes of ligand-receptor interactions and directional communication strength (i.e. connection weight) between cell populations and treatment conditions, we used the R package CellChat v2.2.0 ^122^. “Pallium-4” and “Perithelial” clusters were excluded from the analysis since no specific cell type was assigned and a limited cell number was present. Mouse orthologs of the *L. dimidiatus* were identified as described in the “ortholog gene assignment” section. Then, each dataset (ambient and elevated CO_2_ treatments) was processed separately following a standard pipeline within CellChat workflow by using the “cellChatDB.mouse” and then merged to create the final object. The inference of cell-cell interactions included the computation of the communication probability and identification of significant cellular communication changes between treatments.

## Acknowledgments

We would like to thank all people who gave us suggestions and inputs to make this experiment possible, including Magda Tales from the Oliveira Lab for her inputs on the lab methodology. We are grateful to the members of the Haerty and Macaulay Groups from the Earlham Institute at the Norwich Research Park who never hesitated to help. We also warmly thank Hang Cheong Chan for his precious help with the aquarium facility at SWIMS and Hugo Lee for providing the illustrations of the fish in Figure 1a. We are also grateful to all the members of the Schunter’s Lab for their constant support.

## Author contributions

Conceptualization: C.S., D.D.; Methodology: D.D., C.S.; Investigation: D.D., C.S.; Visualization: D.D. Supervision: C.S.; Writing—original draft: D.D.; Writing—review & editing: C.S., D.D.

## Data and Material availability

All data needed to ensure reproducibility and evaluate the conclusions in the paper are present in the paper and/or the Supplementary Materials. The raw snRNA-seq data generated for this work are deposited on NCBI Gene Expression Omnibus and will be available upon peer review publication. Custom codes and Supplementary Material related to the analyses are also available from GitHub (https://github.com/Deb96-lab/snRNA_brain_ocean_acidification).

