## Supplementary Figures for "Single-nucleus transcriptomics links telencephalic cell signatures to ocean acidification effects on social behaviour in a coral reef fish"

**This PDF file includes:**

Supplementary Figures. S1 to S6

**Other Supplementary Materials for this manuscript include the following:**

Supplementary Excel Tables from. S1 to S19

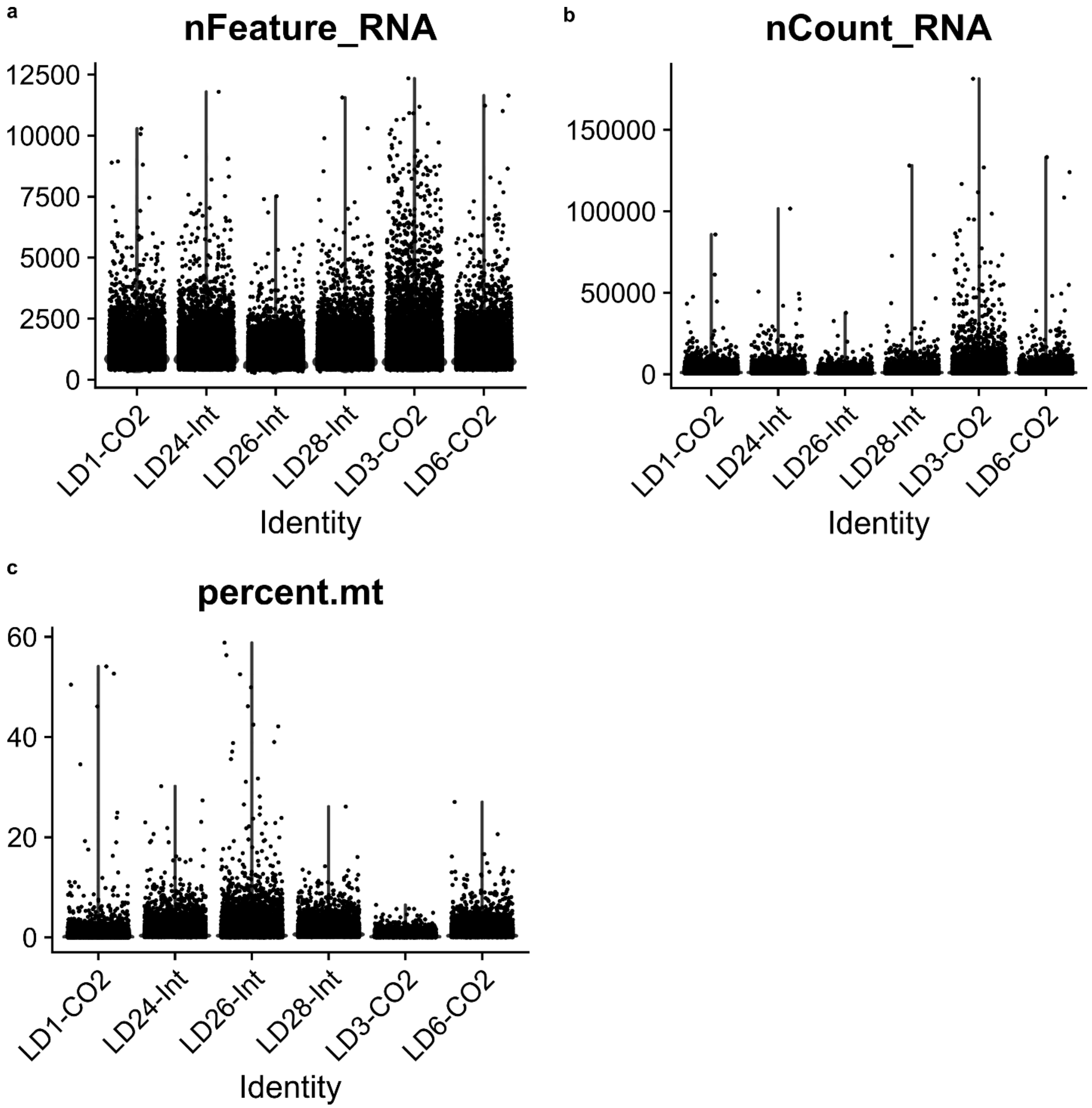


**Fig. S1. Quality control for snRNA-seq before data filtering and data preprocessing.**

(**a**) Number of detected unique genes ("nFeature_RNA"). (**b**) Number of unique molecular identifiers (UMIs, "nCount_RNA"). (**c**) Percentage of mitochondrial genes ("percent.mt"). Data displayed after Cell Ranger pipeline (see the "Single-nuclei RNA sequencing and quality pre-processing" section reported in the Material and Methods for more information). LD24-Int, LD26-Int, LD28-Int are the samples in ambient $CO_2$ treatment, while LD1-CO2, LD3-CO2, LD6-CO2 are the samples in elevated $CO_2$ treatment conditions.

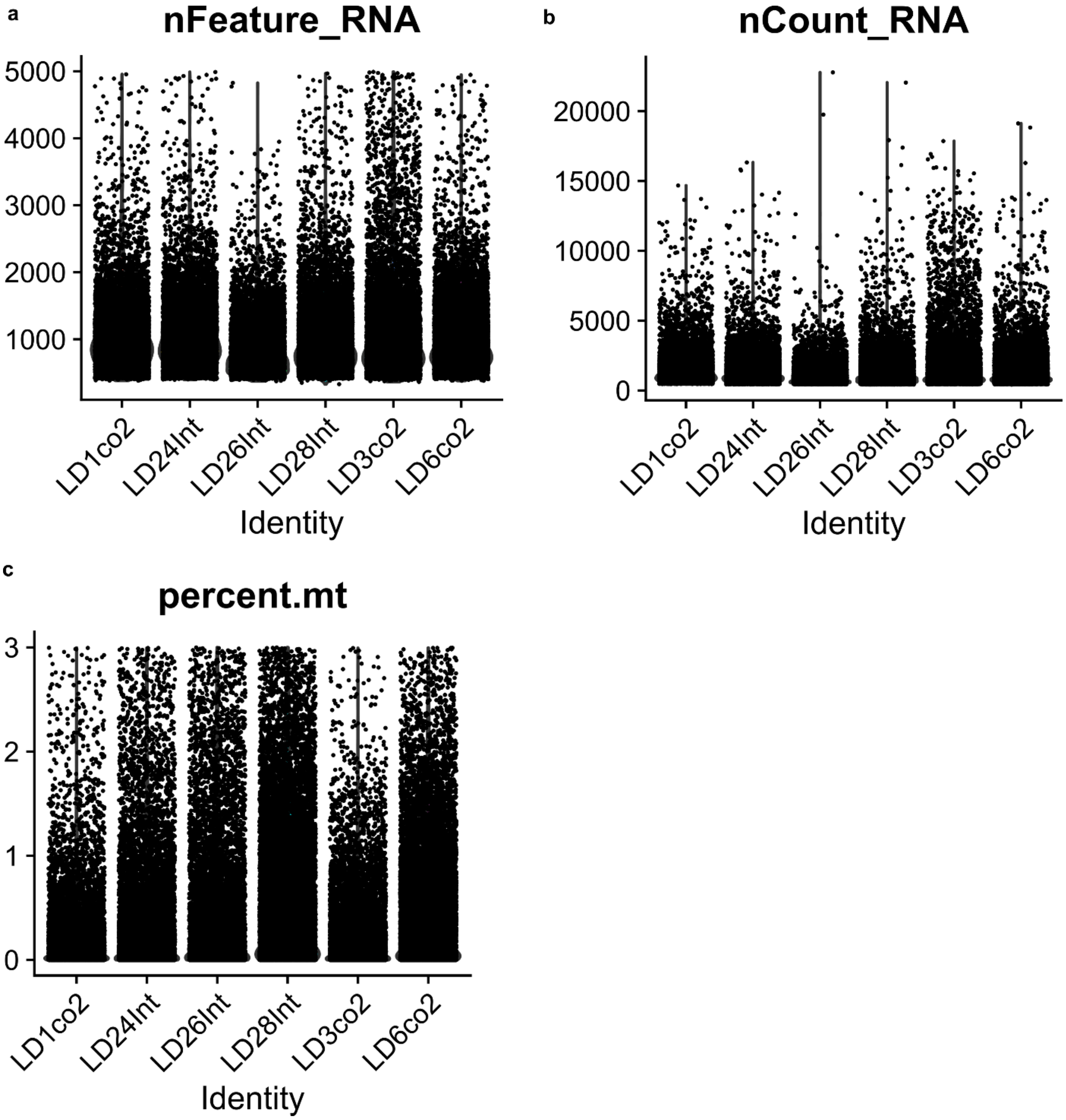


**Fig. S2. Quality control for snRNA-seq after data filtering and data preprocessing.**

(**a**) Number of detected unique genes (“nFeature_RNA”). (**b**) Number of unique molecular identifiers (UMIs, “nCount_RNA”). (**c**) Percentage of mitochondrial genes (“percent.mt”). Data displayed after low-quality cell filtering steps (nFeatures above 250 and below 5000; % of mitochondrial genes < 3; see the “Single-nuclei RNA sequencing and quality pre-processing” section reported in the Material and Methods for more information). LD24Int, LD26Int, LD28Int are the samples in ambient $CO_2$ treatment, while LD1CO2, LD3CO2, LD6CO2 are the samples in elevated $CO_2$ treatment conditions.

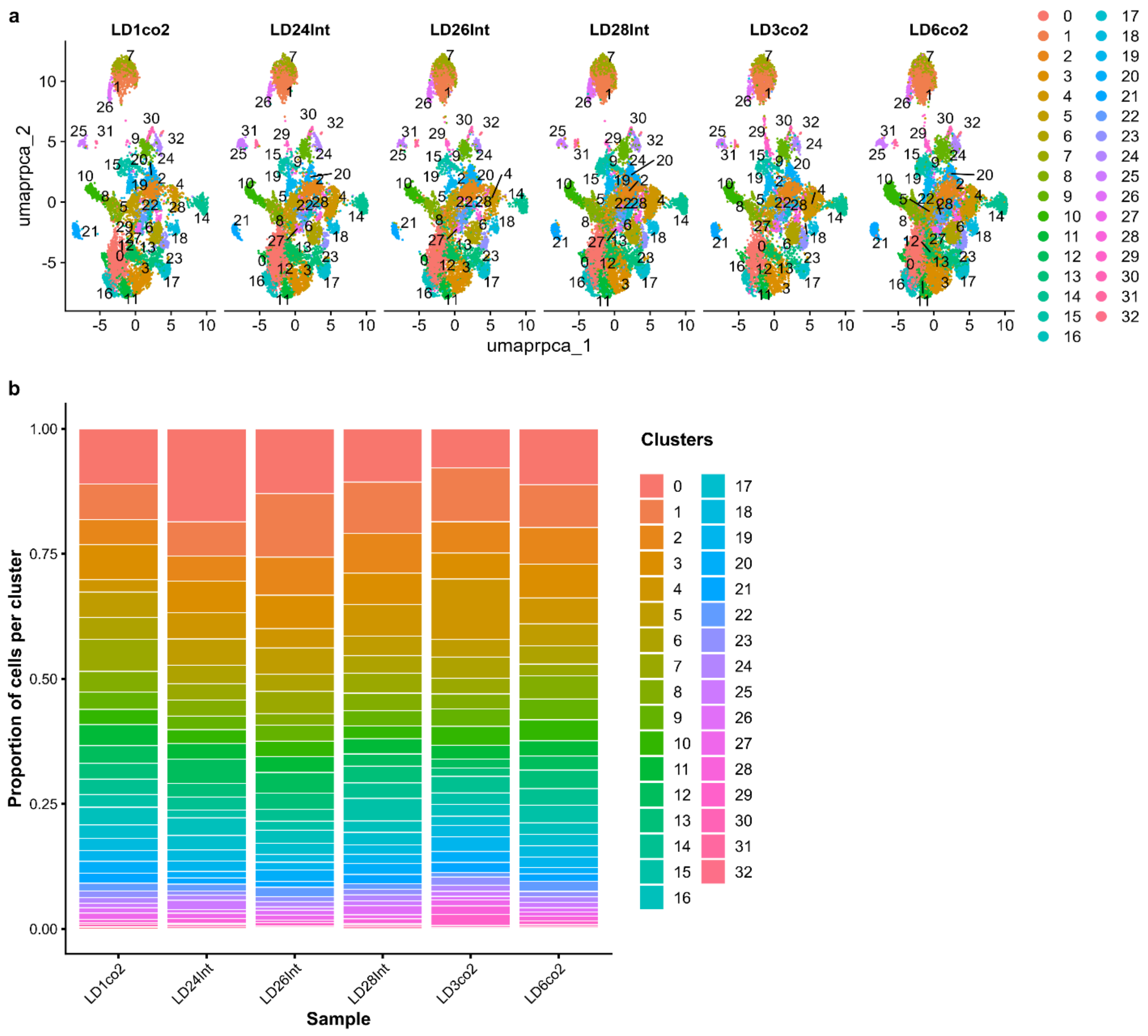


**Fig. S3. Cluster proportions.**

(**a**) UMAP of the cell proportion within each identified cell cluster (*res* = *1.3*). Each treatment is well represented. LD24Int, LD26Int, LD28Int are the samples in ambient $CO_2$ treatment, while LD1CO2, LD3CO2, LD6CO2 are the samples in elevated $CO_2$ treatment conditions. (**b**) Proportion of cells within each identified cell cluster (*res* = *1.3*). LD24Int, LD26Int, LD28Int are the samples in ambient $CO_2$ treatment, while LD1CO2, LD3CO2, LD6CO2 are the samples in elevated $CO_2$ treatment conditions.

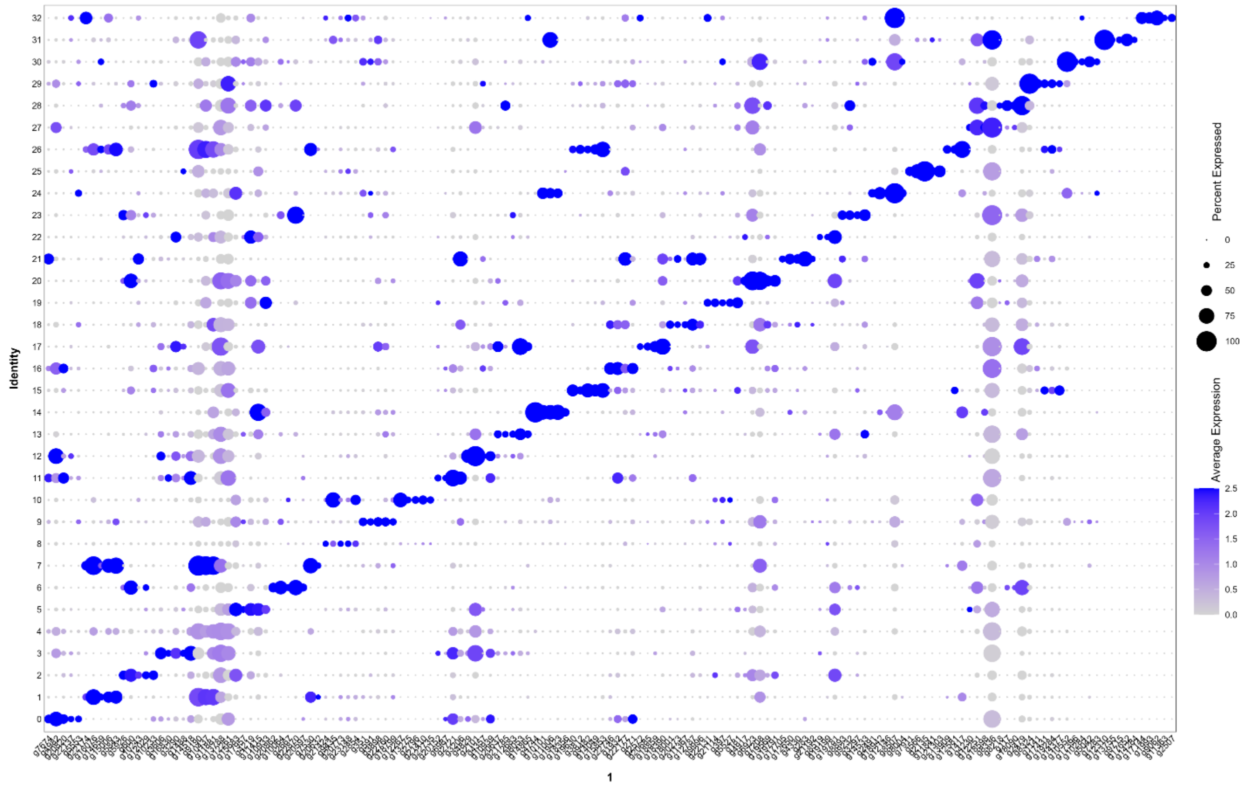


**Fig. S4. Cluster diversity is confirmed by specific gene markers.**

Percentage of nuclei and the average expression of the top 5 marker genes per cluster and cell type family identified from FindAllMarker function. For the corresponding gene labels, see Table S2.

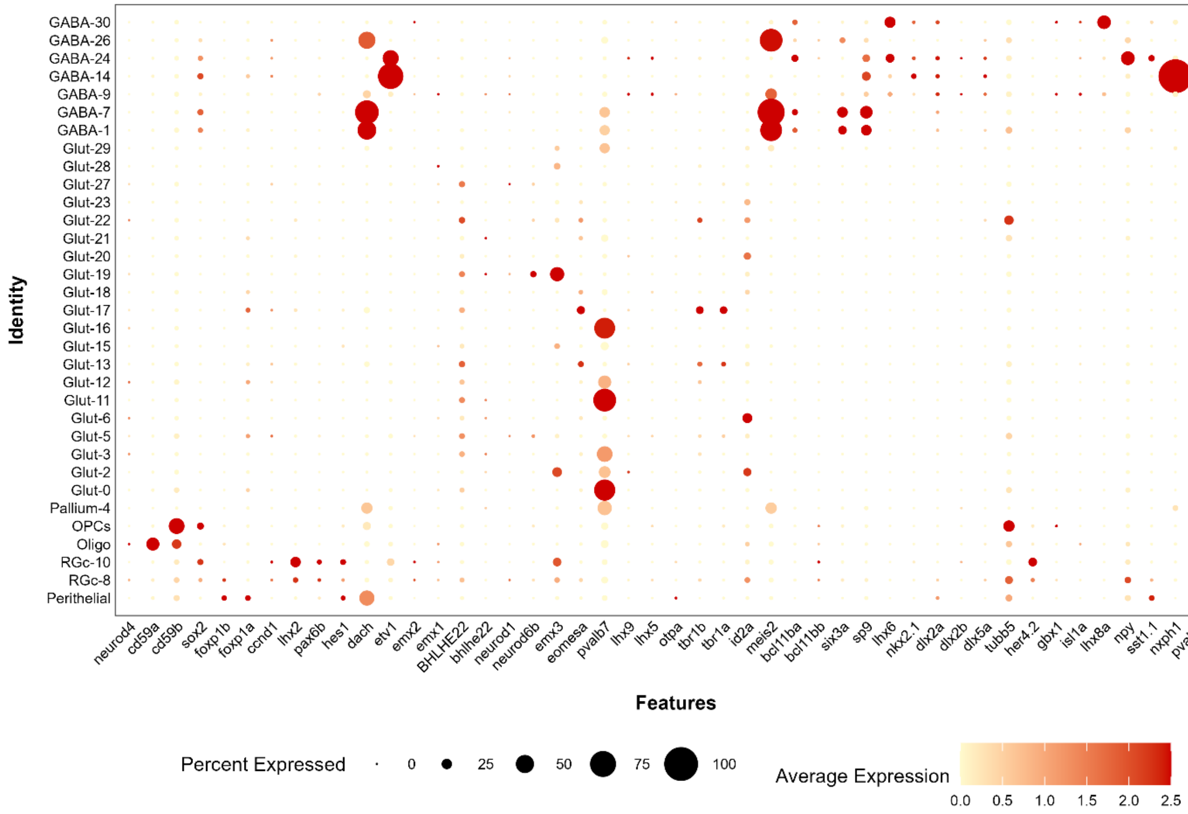


**Fig. S5. Neuroanatomical cell marker genes across *L. dimidiatus* cell clusters.**

Percentage of nuclei and average expression profile of established region-specific neuroanatomical marker genes across all identified cell clusters. Darker colours define higher average expression levels. For more information on the gene marker reference list, see Table S6a.

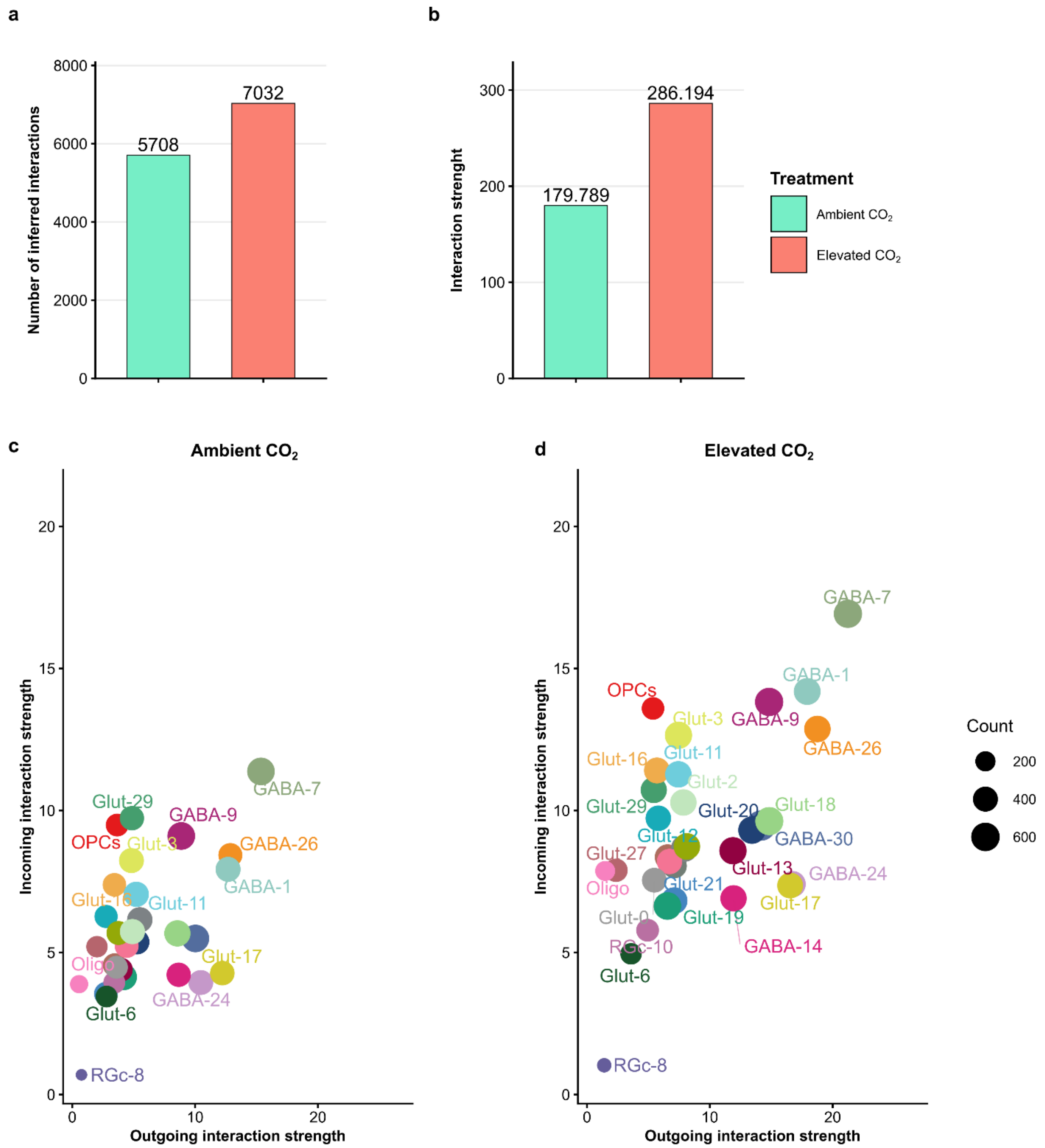


**Fig. S6. Cell-cell communication signalling changes between ambient and elevated $CO_2$ conditions.** Comparison of the total (**a**) number of interactions and (**b**) interaction strength of the CellChat inferred cell-cell communication networks between ambient (light green) and elevated (salmon) $CO_2$ exposure treatments. Number and strength of cellular interactions increased in elevated $CO_2$-treated fish compared to ambient $CO_2$ controls. (**c-d**) Comparison of the outgoing and incoming interaction strength (sending and receiving signals) across the identified cell clusters in (**c**) ambient $CO_2$ and (**d**) elevated $CO_2$ treatment conditions. The overall signal is increased in $CO_2$-treated fish compared to ambient $CO_2$ controls.
